# Stress-dependent growth of breast cancer models arises from a cellular volume checkpoint

**DOI:** 10.1101/2025.07.29.667388

**Authors:** Irish Senthilkumar, Jef Vangheel, Vatsal Kumar, Laoise McNamara, Bart Smeets, Enda Howley, Eoin McEvoy

## Abstract

Mechanoresponsive cell proliferation is a feature of growing tumours, despite the suppression of many other regulatory checkpoints in cancer, but the underlying cell-scale mechanisms driving this behaviour have not yet been established. In this study we propose a biophysical model for cell growth as governed by actively controlled osmolarity, which we integrate with a discrete particle framework to simulate growth and remodelling of breast cancer spheroids. Confinement and biomechanical feedback from the extracellular environment are analysed through a neuralnetwork-accelerated finite element solver. Combining the framework with experiments, our model reveals that stress-dependent spheroid growth can arise from a sizing checkpoint for mitosis. Under sufficient extracellular loading, cell growth is restricted by high hydrostatic forces in competition with osmotic pressure from biomolecule synthesis, which prevents cells from surpassing a critical volume. Our model provides new insight into mechanosensitive growth arrest in breast cancer, potentially serving as a computational tool for analysing growth in a wider range of normal and malignant biological tissues.

## Introduction

Tissue growth is highly regulated by key cell-scale mechanisms, as cells sense mechanical cues and respond to changes in their environment (Di et al., 2023). Non-malignant cells possess several homeostatic controls to cease proliferation and growth under particular conditions. For example, cell cycle progression can be restricted by p53 activation if there is DNA damage or impaired replication (Abuetabh et al., 2022). Cells in dense clusters or monolayers are also subject to contact inhibition of proliferation as mediated by cadherin and Hippo pathway signalling (Gumbiner & Kim, 2014; Puliafito et al., 2012). An early step in cancer initiation is the loss of these such controls, eventually leading to uncontrolled proliferation and metastasis - a characteristic hallmark of cancer. In malignant cells, the loss of contact inhibition increases YAP/TAZ activity, promoting continuous cell proliferation and survivability (Pavel et al., 2018). However, despite the suppression of many cycle checkpoints in cancer, one important regulatory mechanism is preserved across normal and cancerous tissue - mechanical suppression of growth. Proliferation of tumour cells within a restrictive microenvironment leads to the generation of significant stress (Jain et al., 2014; Stylianopoulos et al., 2012). Helmlinger et al., (1997) identified that cell proliferation is suppressed in stiff environments, but interestingly this response is reversible when mechanical stress is relieved. Although such stress-dependence has been widely reported (Curtis et al., 2020; Tan et al., 2024), the underlying biomechanisms are unclear, and a direct link between sub-cellular regulators of cycle progression and macro-scale growth has not yet been established.

In-vitro spheroid models are widely used to model tumour behaviour, providing a platform to study mechanosensitive growth and bridge the gap between the single and multi-cellular scales. Guillaume et al., (2019) demonstrated that in-vitro spheroids can replicate the accumulation of growth-induced solid stress observed in-vivo. Studies have also indirectly captured the spatial distribution of stress within spheroids by tracking the strain of oil droplets (Campàs et al., 2013) and microbeads (Dolega et al., 2017), revealing that cells at the spheroid core experience elevated compressive stress than those at the periphery. By culturing spheroids within microbead-embedded hydrogels, Taubenberger et al., (2019) further showed that spheroid proliferation exerts compressive stress on the microenvironment and proposed that ROCK kinase plays a key role in cell proliferation under mechanically stiff environments. At a cellular level, volume is tightly controlled to ensure regular function and survival. The flow of water across the semi-permeable cell membrane is largely driven by fluctuations in cytosolic ion concentration and osmotic pressure (Reuss, 2012). Cytosolic ion concentration is carefully balanced via a number of active and passive biomechanisms, and disruptions in ion regulation can have severe consequences in downstream tissue-level behaviour. For example, inherited channelopathies arising from ion channel mutations can cause cardiac disorders such as long QT syndrome (Modell & Lehmann, 2006) and anemic disorders such as sickle cell disease (Joiner, 1993). During the cycle, cells increase in size over time. G1 phase growth is typically described by mass accumulation, as in adder models (Nieto et al., 2025), or volume growth, as in sizer models (Facchetti et al., 2017). However, experimental evidence suggests that mammalian cells exhibit hybrid behaviour combining these two paradigms (Cadart et al., 2018). Interestingly, restricting protein synthesis has been shown to reduce volume gain which reinforces that cell growth is driven by the generation of biomolecules (Takauji et al., 2016).

In this study, we develop a novel hydromechanical growth model to describe cell cycle growth. We expand this model to a 3D deformable cell framework for the analysis of discrete multi-cellular spheroid growth. We then couple this framework with a deep neural network-based finite element solver to model the biomechanical behaviour of the ECM and loading induced during growth. Finally, we validate our model predictions with in-vitro growth of breast cancer (T47D and 4T1) spheroids in hydrogels of varying stiffness, revealing the associated mechanisms underlying stress-dependent tumour growth.

## Results

### A hydromechanical model for cell growth and volume control

#### Growth arising from biomolecule synthesis

Tissue growth arises from single-cell growth and remodelling whereby during the cell cycle, cells grow over time and subsequently undergo mitosis. Cell volume growth is largely driven by the synthesis of biomolecules such as proteins, ribosomes and mRNA during the G1 phase (Bernstein et al., 2007; Polymenis & Aramayo, 2015; Riba et al., 2022) (Fig. 1a). Many of these biomolecules are unable to permeate the cell membrane and therefore remain within the cytoplasm (Cooper, 2000; Görlich & Kutay, 1999), increasing the internal osmotic pressure of the cell and subsequently giving rise to water influx; water flows across the semi-permeable cell membrane into a region of high solute concentration (Lopez & Hall, 2023; Zeuthen et al., 2013). Assuming dilute conditions, the osmotic pressure contribution of these impermeable biomolecules for a given cell *i* is derived using the Van’t Hoff equation Π_*x,i*_ = *N*_*x,i*_*RT*/*V*_*i*_, where *N*_*x,i*_ is the impermeable biomolecule concentration, *R* is the gas constant, *T* is the absolute temperature, and *V*_*i*_ is the cell volume. We consider that biomolecules are synthesized at a constant rate *α*_*x*_ such that *dN*_*x,i*_/*dt* = *α*_*x*_ until a saturation value of *N*_*xs*_ is reached, associated with reduced synthesis due to macromolecular crowding (Alric et al., 2022). Synthesis directly leads to cell growth, as the associated increase in osmotic pressure promotes water transport. This water flux across the membrane is described by *J*_*v,i*_ = −*L*_*p,i*_(Δ*P*_*i*_ − ΔΠ_*i*_), where *L*_*p,i*_ is the permeability of the cell membrane, and Δ*P*_*i*_ and ΔΠ_*i*_ are the hydrostatic and osmotic pressure differences across the cell membrane respectively. Cell volume is further controlled by the concentrations of internal and external ion species, which can be transported across the membrane. Considering a number of internal permeable ions *N*_*p,i*_, the total osmotic pressure difference across the cell membrane can be stated as ΔΠ_*i*_ = Π_*x,i*_ + Π_*p,i*_ − Π_*ext*_, where Π_*p,i*_ is the osmotic pressure contribution of the permeable ions and Π_*ext*_ is the osmotic pressure of solutes in the microenvironment. The change in cell volume then follows as *dV*_*i*_/*dt* = *A*_*i*_*J*_*v,i*_, where *A*_*i*_ is the surface area of the cell, leading to:

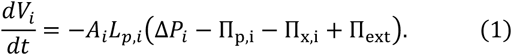

Water influx causes the membrane to stretch as the cell swells in size (Chugh & Paluch, 2018; Kelkar et al., 2020). Cortical stress can be written as *σ*_*i*_ = *σ*_*p,i*_ + *σ*_*a,i*_, where *σ*_*p,i*_ is the passive stress arising from the deformation of the actin network and *σ*_*a,i*_ the active cortical stress. Mechanical force balance for a single spherical cell of radius *r*_*i*_ dictates that the cortical tension can be related to the hydrostatic pressure difference across the membrane Δ*P*_*i*_ = *P*_*i*_ − *P*_*ext*_, where *P*_*i*_ and *P*_*ext*_ are the internal and external hydrostatic fluid pressures respectively. If we consider that cells may also experience further external mechanical loading, for example due to contact stress *σ*_*ext,i*_, the cortical stress may also be written as *σ*_*i*_ = (Δ*P*_*i*_ − *σ*_*ext,i*_)*r*_*i*_/2*h*_*i*_, where *h*_*i*_ is the combined thickness of the cortex and membrane.

**Figure 1:**
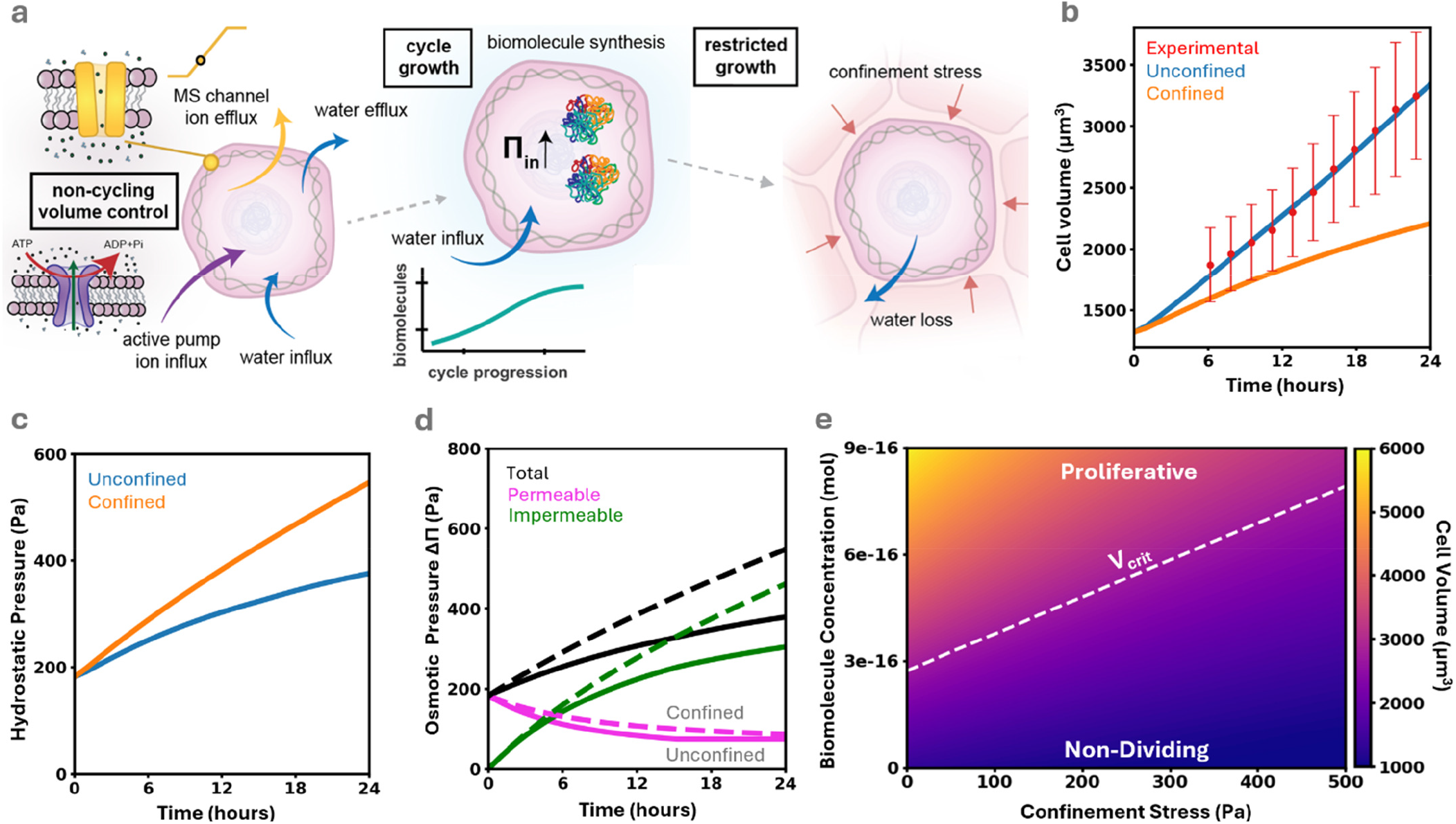
Influence of confinement on single-cell growth. **(a)** Hydrostatic and osmotic pressure gradients allow cells to exchange water and solutes across the semi-permeable membrane through the synthesis of impermeable biomolecules. Contact stress arising from cell loading increases hydrostatic pressure and causes fluid loss, preventing cells from reaching the critical mitotic volume. **(b)** The cellular growth curve was calibrated to align with single-cell growth (mean ± SD) of the HeLa cell line (Cadart et al., 2022). Cell confinement inhibits volume growth. **(c)** Contact stress also increases intracellular hydrostatic pressure. **(d)** Biomolecule synthesis is the primary contributor to the change in intracellular osmotic pressure. Contact stress drives water efflux and reduces cell volume, in turn increasing osmotic pressure. **(e)** A competition between hydrostatic and osmotic pressures, induced by the balance between contact stress and biomolecule synthesis, governs cell cycle growth under mechanical loading.

#### Fluid and ion exchange with the extracellular environment

As mentioned, in addition to impermeable biomolecule synthesis, cell volume is further dependent on the concentration of permeable ions *N*_*p,i*_ within the cytoplasm. Following from previous work (Jiang & Sun, 2013; McEvoy et al., 2020), the transport of ions across the membrane is facilitated through several passive and active biomechanisms. We assume the existence of a single permeable ion species and ignore the influence of electroneutrality and membrane potential. Cell volume growth increases cortical stress (Liu et al., 2023), in turn opening passive mechanosensitive ion channels (Kung, 2005; Patel et al., 2015) and permitting ion transport in accordance with a concentration gradient at a rate 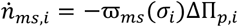, where ϖ_*ms*_ is the channel permeability and *σ*_*i*_ is the cortical stress. The channel permeability is dependent on cortical stress and has been reported to follow a Boltzmann function (Sukharev et al., 1999). In addition to mechanosensitive channels, continuously functional passive leak channels on the cell membrane facilitate ion transport towards the concentration gradient (Alberts et al., 2002) with channel ion flux defined by 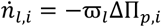, where ϖ_*l*_ is the leak channel permeability. While these passive biomechanisms permit ion flux in the direction of the concentration gradient, active biomechanisms permit ion flux against the concentration gradient. Active ion pumps hydrolyse ATP to overcome the energetic barrier (Kjelstrup & Lervik, 2021) with flux defined by 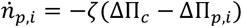 where ζ is the pump rate constant and ΔΠ_*c*_ is the critical osmotic pressure difference above which ATP hydrolysis would not provide sufficient energy for pumping. After taking these passive and active biomechanisms for ion transport into consideration, the internal permeable ion concentration is then determined as 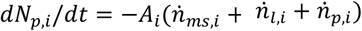, or:

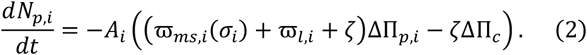

### Impermeable biomolecule synthesis drives cell growth

We first consider the behaviour of our hydromechanical model during growth of individual HeLa cells. Our model suggests that the synthesis of impermeable biomolecules gives rise to an increase in cytosolic osmotic pressure, driving volume growth due to an influx of water through the semi-permeable membrane during the G1 phase (Fig. 1b). The predictions from our model agree with experimental trends of suspended HeLa cells (Cadart et al., 2022), with cell volume increasing almost two-fold from early G1 phase to the start of the M phase. The influx of water into the cytosol elevates intracellular hydrostatic pressure, predicted to increase from 210 Pa at the start of the G1 phase to 340 Pa at the start of the M phase (Fig. 1c), aligning with experimental data of HeLa growth (Fischer-Friedrich et al., 2014). Increases in cell size raise cortical stress as the membrane stretches to accommodate the influx of water, subsequently increasing ion loss by increasing permeability of mechanosensitive ion channels. However, biomolecule synthesis and ion pumping offset associated changes in osmotic pressure, sustaining cell growth (Fig. S1). During growth, cells are subject to a variety of mechanical loads arising from confinement and interactions with the ECM and neighbouring cells. Contact stresses increase intracellular hydrostatic pressure which, in competition with the osmotic pressure, may lead to fluid loss from the cell and a reduction in size (Fig. 1c-d). In turn, reduction of cell volume and cortex area reduces cortical stress and mechanosensitive channel permeability, mitigating excessive reductions in osmotic pressure. Our analysis indicates that growth in confined cells during their cycle is dependent on a competition between the number of synthesized biomolecules and the external loading imposed on the cell surface (Fig. 1e). Bridging from growth to mitosis, Varsano et al., (2017) identified that the likelihood of cancer cell proliferation increases with cell size, facilitating identification of a characteristic volume for cycle progression. Our model suggests that a critical level of stress from confinement may restrict cell growth below such a mitotic volume checkpoint to arrest proliferation (Fig. 1f).

### An agent-based hydromechanical cell model for multicellular growth

Having explored the influence of mechanical loading on single cell growth, we extend our hydromechanical model to analyse the growth of multi-cellular tumour spheroids. We integrate our framework with a 3D deformable cell model, where tumour tissue is described as an active cellular foam (Cuvelier et al., 2023; Ongenae et al., 2024; Vangheel et al., 2025). Individual cells are characterized by a thin viscous shell with viscosity *η*_*c*_ and surface tension *γ* that represents the cortex actomyosin remodeling, and contraction, respectively. The cell can interact with neighboring cells by adhesive interactions parameterized by adhesive tension *ω* that together with cortex tension determines the cell-cell contact angle (Fig. 2a). Extracellular interactions are modelled by repulsive pressures. Furthermore, cell-cell interactions include wet friction with coefficient ξ. The cortex shell is discretized by a triangulated surface mesh where nodal positions act as the relevant degrees of freedom. A detailed description of this deformable cell model can be found in Supplementary Note 2. The cellular environment is assumed to be overdamped, allowing for inertial forces to be neglected (Purcell, 1977). Cell volume growth is described as outlined in eqns 1-2, with predicted volume achieved within discrete cells via a proportional integrative pressure controller (see Methods). We consider that cycle progression and mitosis is subject to a volume checkpoint (see Methods) as highlighted in Fig. 1. Cytokinesis is simulated by constricting a central ring of nodes on the cell surface, symmetrically dividing fluid volume, permeable ions, and impermeable biomolecules between daughter cells.

**Figure 2:**
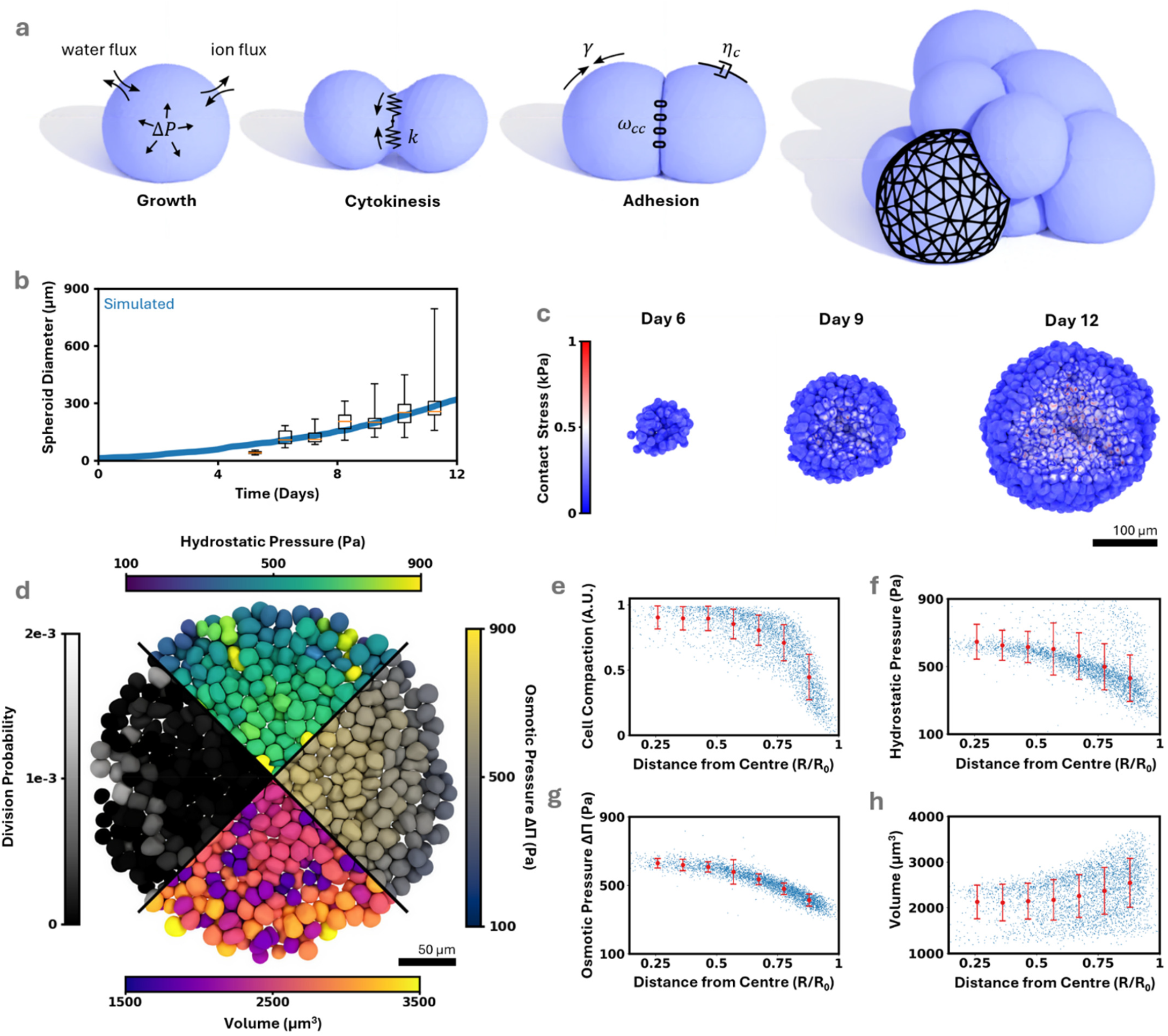
Spatial Variations in cell volume arise from mechanical loading in unconfined HeLa spheroid growth. **(a)** The viscous deformable cell model is described by active foam theory. **(b)** Spheroid growth curve aligns with experimental data (Jang et al., 2024). **(c)** A volume cut-out exposes the inner composition of the spheroid over time. Proliferation generates contact stress due to cell-cell adhesive forces. **(d)** A section cut on the equatorial plane uncovers spatial variations in cell-level observables. **(e)** Increased contact stress at the core leads to greater compaction. The competition for core cell volume regulation between **(f)** hydrostatic pressure - induced by contact stress, and **(g)** osmotic pressure - induced by biomolecule synthesis, favours hydrostatic pressure at the core, leading to volume efflux and **(h)** subsequent cell volume loss. All error bars show mean ± SD.

We proceed to investigate the role of confinement on spheroid proliferation and growth inhibition. Our combined agent-based hydromechanical deformable cell model predicts that cells increase in size through biomolecule synthesis and increasingly undergo mitosis when their volume reaches a characteristic mitotic volume *V*_*crit*_. Cells proliferate over time, leading to an increase in spheroid size, with cells bound together by cadherin-based adhesion to mediate spheroid compaction. We achieve close agreement between simulated spheroid diameter and experimental observations of HeLa spheroid growth (Jang et al., 2024) to verify model predictions (Fig. 2b). Notably, there is an accumulation of contact stress within proliferating spheroids, prominent in the spheroid core (Fig. 2c). This spatial distribution in contact stress has been observed experimentally (Nia et al., 2016), and gives rise to spatially heterogeneous cell behaviour (Fig. 2d). Cell compaction, defined here as the ratio between a cell’s in-contact and total surface area, is higher at the core and tends lower towards the periphery (Fig. 2e). This spatial variance in cell compaction suggests that cells at the core undergo significant deformation, whereas exposed peripheral cells largely maintain a spherical morphology. Our model predicts a reduction in ion efflux at the core due to the reduction in exposed cell surface area arising from increased cell-cell contact (Fig. S2a-b). As with the single-cell model, contact stress leads to an increase in cytosolic hydrostatic pressure (Fig. 2d, 2f) and osmotic pressure (Fig. 2d, 2g). As cell volume growth is regulated by the competition between hydrostatic and osmotic pressure, the spatial imbalance between these pressures drives a spatial variance in cell volume. Hydrostatic pressure exceeds osmotic pressure at the core, leading to volume loss and growth inhibition as cells cannot surpass the volume checkpoint. Conversely, osmotic pressure is higher at the periphery, driving cell growth and proliferation (Fig. 2d, 2h), reflecting experimental trends (Browning et al., 2021).

### Neural network-driven finite element solver to model the extra-cellular matrix

Importantly, most solid tumours are not freely suspended and instead they are encapsulated within extra-cellular matrix (ECM) proteins such as collagen and fibrin (McKee et al., 2019). Complex mechanical and biochemical feedback between the cells and such ECM influences growth and may guide tumour progression (Humphrey et al., 2014; Yoshii et al., 2016). To simulate contact between cells and matrix, we developed a deep neural-network–accelerated finite element (DNN-FE) solver. The DNN is designed to predict the displacements of ECM surface nodes (see Methods) for a given load distribution arising from contacting cells (Fig. 3b). Synthetic data (6,000 samples) were generated for training and validation processes using the finite element software, with the biomechanical behaviour of the ECM described by a neo-Hookean constitutive law. The DNN demonstrates stable convergence of the trainable parameters, as evidenced by the decreasing training and validation loss curves (Fig. 3c). Details of the training data generation pipeline and model hyperparameters are provided in the Methods and Supplementary Note 4. Our DNN-FE framework accurately predicts the deformation of the ECM surface, achieving 99.5% prediction accuracy. The average mean absolute error between the ground truth and predictions is ~3 µm (Fig. 3d), indicating excellent agreement with the Abaqus finite element software for a range of loading conditions (Fig. 3e-f). Crucially, arising from the low computational cost associated with neural network predictions, our framework improves computational efficiency 500,000-fold relative to equivalent CPU-based finite element solvers (Fig. 3g).

**Figure 3:**
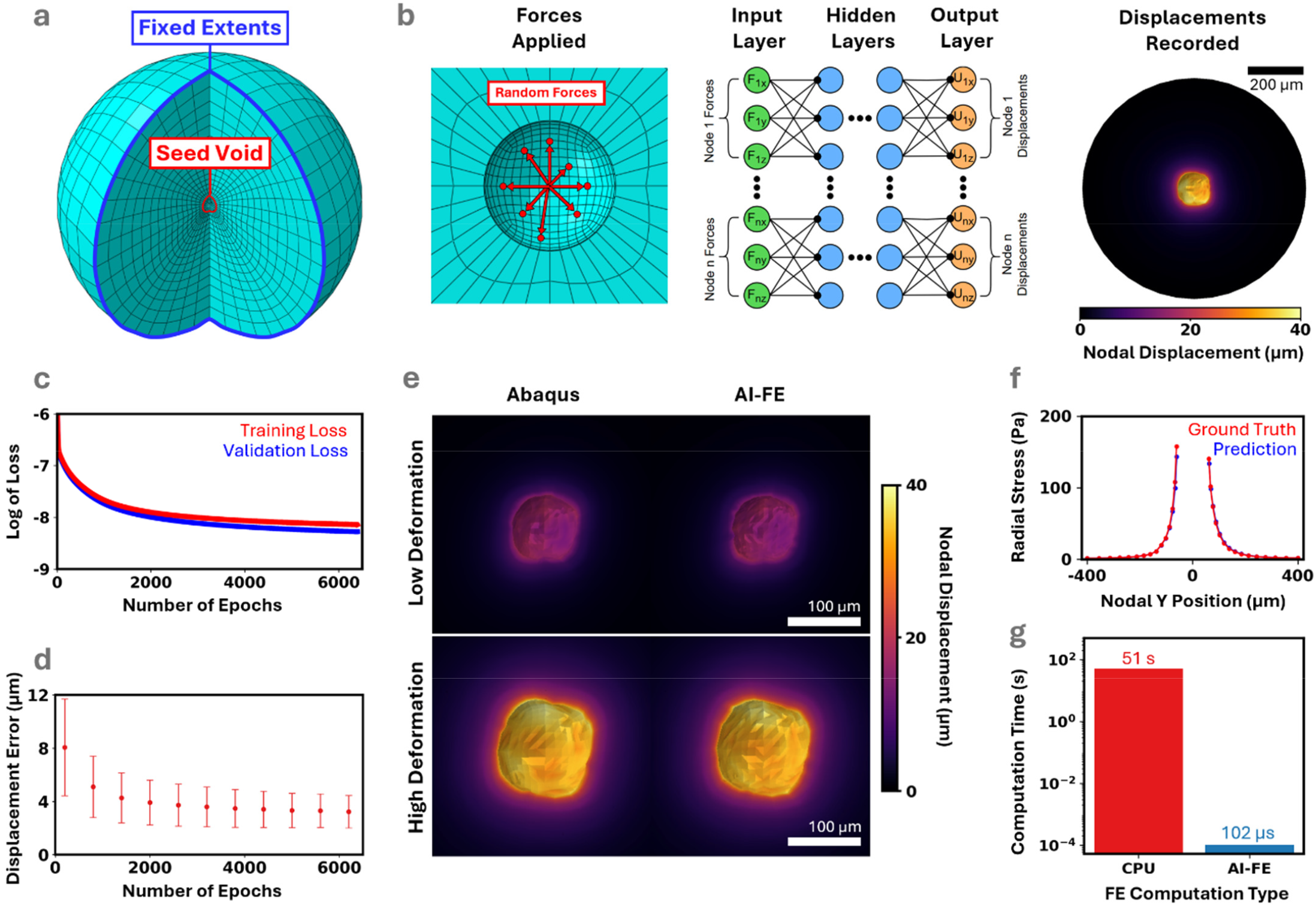
Neural network-driven finite element solver to model the mechanical response of the matrix. **(a)** The matrix is represented as a solid sphere with a central seeding void. The outer nodes are fixed in position. **(b)** Forces applied on the seeding surface deforms the surrounding matrix. The Cartesian components of these applied nodal forces serve as inputs to the DNN, which outputs the corresponding Cartesian nodal displacements. **(c)** As the DNN trains over multiple epochs, the training loss, and consequently **(d)** the error between predicted and true nodal displacements, gradually decreases (mean ± SD). **(e)** The DNN accurately predicts the displacement of the seeding surface nodes with high accuracy for a wide range of deformations. **(f)** Predicted radial stress along the Y axis nodal path closely aligns with the ground truth. **(g)** The DNN-FE framework predicts FE analysis 500,000 times faster than traditional CPU-based FE methods.

### Cell proliferation rates reduce in spheroids within stiff ECM

Expanding tumour spheroids compress and stretch surrounding ECM, generating normal and tangential stress. To characterise these interactions within our computational framework, a linear force contact model generates repulsive forces between cells and the ECM surface (see Methods). Rapid convergence of the contact model is achieved through a viscous soft relaxation algorithm. This method for contact convergence allows consistent contact to be maintained under high deformations, verified by simulating microbead indentation of the ECM surface (Fig. 4a). Predictions from the combined cell-matrix model indicate that ECM deformation and stress are predominantly localised adjacent to the spheroid surface and increase significantly during growth (Fig 4b). Radial and circumferential stress in the ECM constrains the spheroid, elevating cell-cell contact stresses (Fig. 4b). This leads to high levels of compaction and cell deformation within the spheroid. Under this constraint and at this scale, contact stresses, hydrostatic pressure and osmotic pressure are predicted to be spatially uniform (Fig. 4c, e, f) compared to freely suspended spheroids (Fig. 2), thereby minimizing cell volume variability.

**Figure 4:**
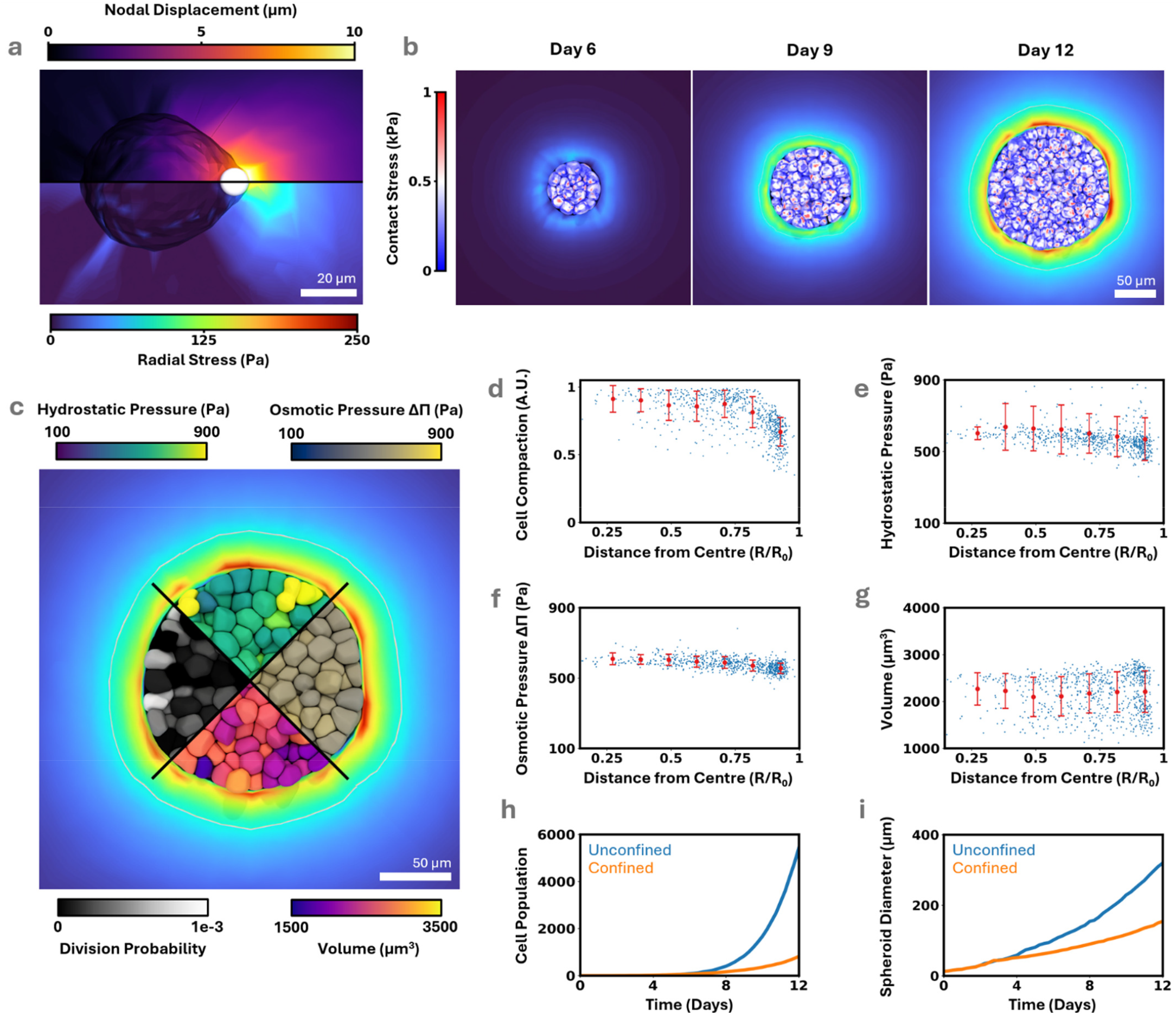
Combined model simulates HeLa spheroid growth in matrix. **(a)** Microbead indentation against the seeding surface highlights contact model stability under high deformations. **(b)** Section cut of confined spheroid growth over time reveals uniform loading across the spheroid, with radial stress increasing alongside proliferation and deformation. **(c)** Section cut of confined spheroid growth shows minimal spatial variance in cell-level observables. **(d)** Homogeneity in contact stress indicates uniform loading across the spheroid, leading to uniform hydrostatic pressure and **(f)** osmotic pressure, ultimately **(g)** eliminating the spatial variance in cell volume. Spheroid growth under ECM confinement suppresses proliferation, leading to a significant decrease in **(h)** cell population and **(i)** spheroid diameter in comparison to unconfined spheroid growth. All error bars show mean ± SD.

In a direct comparison with unconfined spheroid growth, mechanically-induced stress from the surrounding ECM significantly reduces cell proliferation. Consequently, the overall cell population within the spheroid is reduced (Fig. 4h), leading to a significant reduction in spheroid diameter when compared to unconfined growth (Fig. 4i).

### Experimental validation of stress-dependent T47D spheroid growth

To assess the predictive capabilities of our combined model, we expanded human breast cancer cells (T47D) in media and seeded them at low density in gelatin hydrogels. Initial Young’s Moduli of hydrogels were established at 0.58 kPa, 0.85 kPa, and 1.1 kPa. Spheroids spontaneously formed and grew over 12 days, with media changed every two days. Constructs were fixed, permeabilized, and stained for actin (phalloidin) and nuclei (DAPI), with Z-stack images obtained from confocal microscopy. Our computational framework suggests that reducing hydrogel stiffness from 1.1kPa to 0.58kPa leads to a 50% increase in spheroid diameter following 12 days of growth, in direct agreement with our experimental data. Tumour cell proliferation increasingly deforms the surrounding hydrogels over time, with a corresponding increase in pressure applied to the spheroid surface. Notably, the peak radial stress induced by proliferation was similar in all three cases (~400 Pa, Fig. 5a), indicating a maximum growth-induced stress than can be exerted by T47D spheroids. Predictions from our model are also consistent with experimentally-recorded spheroid diameter at all three timepoints (Fig. 5b), spheroid population (Fig. 5c) and average cell volume (Fig. S3). Our simulations suggest that spheroids in stiffer hydrogels experience elevated levels of cell compaction at an equivalent size, with cell compaction increasing towards the core (Fig. 5d). Mean cytosolic hydrostatic pressure is predicted to increase with increasing hydrogel stiffness (Fig. 5e). This, in turn, reduces the rate of cell growth and restricts cells from surpassing the mitotic checkpoint, limiting spheroid expansion. We further validated out model against growth of 4T1 spheroids (Fig. S4), identifying that the underlying mechanism of stress-dependent growth appears to be conserved.

**Figure 5:**
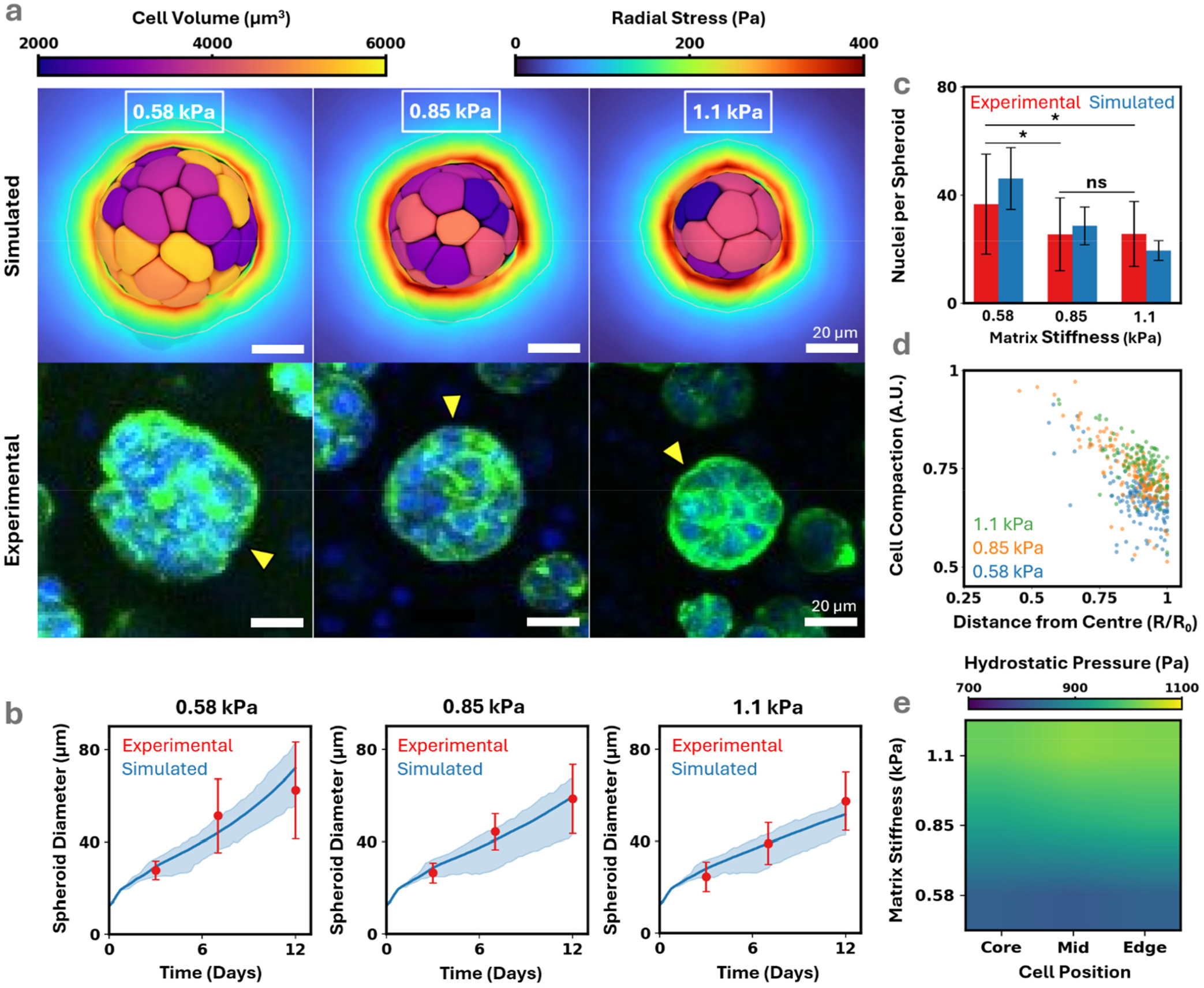
T47D spheroid growth in matrices of varying stiffness. **(a)** Section cut of confined growth for three different hydrogel stiffnesses compared with representative experimental images. **(b)** Predicted confined spheroid growth curves closely aligns with experimental observations. The shaded blue region representing the minimum and maximum spheroid diameters predicted across simulations. **(c)** Predicted number of cells per spheroid closely aligns with experimentally measured nuclei per spheroid. **(d)** Simulations predict an increase in cell compaction with increasing hydrogel stiffness (day 12), tending higher towards the spheroid core. Measurements taken at a 50µm diameter in all cases. **(e)** Hydrostatic pressure values (day 12) are spatially consistent within the spheroid, increasing with hydrogel stiffness. All error bars show mean ± SD. Significance compared using two-way ANOVA with a Bonferonni’s post hoc test with *p≤0.05.

## Discussion

Although mechanical stress has long been implicated in tumour growth, the precise subcellular mechanisms by which mechanical forces regulate cell cycle progression remain poorly understood. In this study, we propose that hydromechanical regulation of cell volume - driven by a competition between osmotic pressure from biomolecule synthesis and hydrostatic pressure induced by mechanical loading - forms a critical link between micro-scale cell behaviour and macro-scale tissue growth.

Cell volume is tightly regulated by numerous underlying biomechanisms including mechanosensitive ion channels, active ion pumps, and leak channels that actively collaborate to manipulate intracellular osmolarity and water flux through the cell membrane. Initially considering a single cell system, mechanical loading promotes water efflux and reduces cortical tension, thereby lowering mechanosensitive channel permeability and ion loss to the microenvironment. Meanwhile, the synthesis of impermeable biomolecules and active ion pumps drive an increase in cytosolic ion concentration and osmotic pressure. Our model reveals that mechanical stress arising from confinement suppresses cell growth not merely by external compression, but by actively disrupting the balance between intracellular hydrostatic and osmotic pressure for growth. Confinement induces a net efflux of water, inhibiting volume growth and prevents cells from reaching the mitotic volume checkpoint for division. While continuum (Carrasco-Mantis et al., 2023) and agent-based (Ghaffarizadeh et al., 2018; Mirams et al., 2013) models have advanced our understanding of spheroid growth, existing frameworks lack the ability to resolve subcellular mechanical phenomena for linking mechanical stress to cell cycle regulation. Continuum models cannot capture the cell-level response leading to heterogeneous stress states, and traditional agent-based models treat cells as rigid spheres, limiting their ability to capture the morphological evolution of tumour cells and mechanical signals transmitted by the microenvironment. Arising from rapid advancement in hardware acceleration and software optimisation, recent studies have advanced a new class of agent-based models that incorporate deformable cells to accurately capture the mechanics of cell-cell interactions (Ongenae et al., 2024; Van Liedekerke et al., 2019). Building on this progress, we integrate our hydromechanical model with a deformable cell framework, allowing us to directly link subcellular osmotic dynamics to macroscopic tissue growth during multicellular proliferation.

We find that mechanical loading arising solely from cell-cell mechanical interactions is sufficient to generate spatial heterogeneity in contact stress and growth. As cells proliferate, they exert mechanical forces onto their neighbours, leading to the development of solid stresses and hydrostatic pressure particularly within the spheroid core (Dolega et al., 2017; Nia et al., 2016). Our simulations predict that this spatial distribution of stress disrupts the balance between intracellular hydrostatic and osmotic pressures in a spatially-dependent manner, with core cells experiencing suppressed growth and reduced proliferation. This heterogeneity in cell volume onset by mechanical loading offers a potential explanation for the characteristic rim-core proliferation patterns observed in spheroids in-vitro (Montel et al., 2012). Cancer cells are typically embedded within a confining ECM and as cells proliferate, they deform the surrounding ECM (Fig. 6a). Although the mechanical interplay between cells and the ECM has been known to guide tumour growth and progression (Helmlinger et al., 1997; Kumar et al., 2024), the role of the ECM is often overlooked by current in-silico models. To address this gap, we next extended our framework to examine how matrix confinement alters multicellular growth and proliferation. We present a computational framework that combines neural networks and finite element methods to simulate matrix deformation in response to cell-matrix interactions with high spatial accuracy and computational efficiency, providing a viable replacement to CPU-based finite element solvers. Unlike freely-suspended spheroid growth where stress gradients gradually arise from cell-cell interactions, the presence of a mechanically resistive ECM imposes a uniform stress gradient across the spheroid. This spatial homogeneity in mechanical loading inhibits cell volume and largely eliminates spatial variations in proliferation. Validated against in-vitro spheroid growth of the T47D and 4T1 cell lines, our combined model captures the relationship between ECM stiffness and proliferation, reinforcing the capability of our hydromechanical cell-ECM model for understanding stress-dependent growth as regulated by matrix stiffness and a critical volume checkpoint (Fig. 6b).

**Figure 6:**
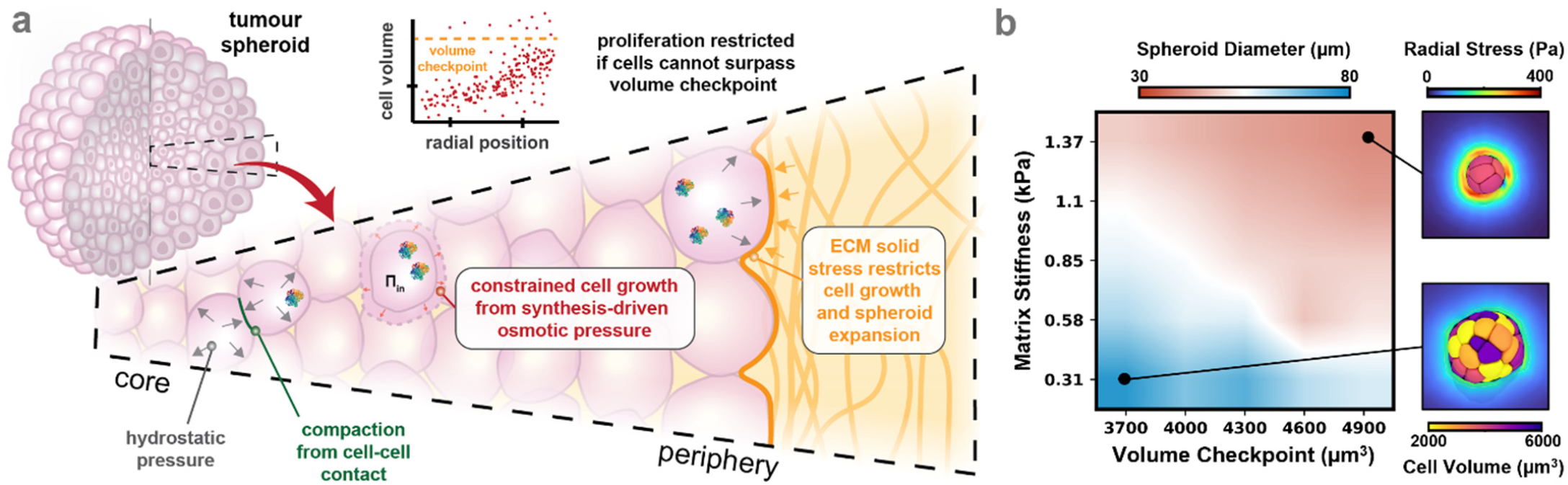
Matrix stiffness and critical mitotic volume are key parameters in cell proliferation under matrix confinement. **(a)** Confinement arising from cell-cell and cell-matrix mechanical interactions inhibits proliferation by restricting cell growth below the volume checkpoint for division. **(b)** An increase in matrix stiffness volume leads to an increase in contact stress and volume loss during proliferation. An increase in critical mitotic volume prevents increases the volume required for mitosis.

These findings, particularly the suppression of proliferation in stiff ECMs, align with a growing body of experimental evidence linking mechanical stress to cell cycle arrest across both single-cell and tissue scales. Huang et al., (1998) demonstrated that increased cytoskeletal tension in endothelial cells promotes G1 arrest through p27^Kip1^ kinase upregulation implicating mechanical stress as a barrier to proliferation. More recently, studies by Delarue et al., (2014) and Taubenberger et al., (2019) revealed spatial patterns of cell cycle arrest in multicellular spheroids exposed to mechano-osmotic stress or confinement within the ECM. Their studies may partly explain how a critical volume checkpoint is underpinned by either the dilution of cell cycle inhibitors (CDKI, RB) (Petermann et al., 2002; Yoon et al., 2012; Zatulovskiy et al., 2020), or macromolecular crowding (Alric et al., 2022). Our work identifies that mechanical stress interrupts the balance between intracellular hydrostatic and osmotic pressure, inducing water efflux and reducing cell volume below the critical mitotic threshold. This hydromechanical imbalance, validated against single-cell and multi-cellular growth, provides a unified explanation for stress-sensitive proliferation and connects cell cycle control to tissue-level mechanics. The suppression of proliferation under mechanical stress has important consequences for cancer treatment, particularly for chemotherapies that rely on active cell cycling. It has been established that mechanical stress inhibits the efficacy of therapeutic drugs by compressing vasculature (Neophytou et al., 2025) and limiting perfusion (Provenzano et al., 2012), and recent research by Rizzuti et al., (2020) further suggests that growth-induced mechanical stress also disrupts the efficacy of cell cycle-dependent drugs such as gemcitabine by inducing cell cycle arrest. Beyond its relevance to tumour biology, the integration of subcellular hydrodynamics into tissue-scale models opens new opportunities to explore how stress regulates cell fate in diverse contexts. Further studies could leverage this computational framework to analyse the role of mechanical stress and hydromechanical regulation in other physiological processes such as wound healing (Agha et al., 2011; Tanner et al., 2009) and morphogenesis (Dasgupta et al., 2018), providing a foundation to explore how mechanical forces shape cellular dynamics across diverse biological systems.

## Materials and methods

### Single cell simulation procedure

For the single-cell analyses, the hydromechanical model was implemented as a system of discretized ordinary differential equations in C++. Richardson extrapolation was used to improve solution accuracy. Initial conditions were identified by solving the equations at steady state. Cell behaviour was simulated over 24 hours. In the confined single-cell case, a time-dependent uniform surface pressure *σ*_*ext*_ was applied on the cell surface, increasing to a peak stress of 250 Pa applied at 24 hours (Fig. S1a). The motivations for all cell parameters are presented in Supplementary Note 3.

### Deformable cell framework

The tumour tissue is modelled as a foam, with details provided in Supplementary Note 2. Briefly, single cells are represented as pressurized bubbles assuming the single cell shape is governed by the actomyosin cortex that can be represented as a viscous fluid on long timescales. The cortex contractility is then modelled with a surface tension *γ*_*c*_ and the cortex remodelling is described as an effective 2D viscosity *η*_*c*_. The in-plane cortex tension can be written as ***σ***_2*D*_ = *η*_*c*_Av + *γ*_*c*_**I**, with **v** = (u, v) the in-plane velocity of the cortex. The divergence of tension is balanced by viscous frictional forces with neighboring cells/ECM and medium with viscosity *η*_*f*_ as A ⋅ ***σ***_2*D*_ = *ξ*^∥^(Δv) + *η*_*f*_v, where *ξ*^∥^ is the tangential friction constant and Δv the relative velocity with neighbouring surfaces. For curved surfaces, the cortex tension generates out of plane forces that is balanced by hydrostatic Δ*P*_*h*_ and contact pressure *P*_*c*_, and medium viscous friction as 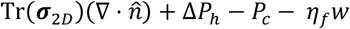, with 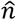 the normal of the surface pointing outwards, and w the outwards surface velocity. We consider that cycle progression and mitosis is subject to a volume checkpoint (Varsano et al., 2017), with a probability defined by ρ_*d,i*_(*V*_*i*_) = *λ*/ (1 + *exp*(−ϱ(*V*_*i*_/ *V*_*crit*_))), where λ is a tunable parameter representing the proliferative capacity of the current cell line, ϱ is a parameter describing the stochastic nature of mitosis and *V*_*crit*_ is the critical mitotic volume. Cell compaction is measured as a ratio of a cell’s surface area in contact to its total surface area.

### Deep neural network-driven finite element analysis framework

The ECM is represented as a solid spherical geometry of outer radius 400 µm with a central spherical seeding void of inner radius 25 µm. For T47D and 4T1 analyses, the outer and inner radii are 200 µm and 15 µm respectively. The outer mesh nodes are fixed in position to prevent displacement and rotation. Using the Abaqus finite element software (version 2021), the ECM part is meshed with 19,440 linear tetrahedral elements (C3D4). A neo-Hookean material model is used to simulate strain stiffening material of the matrix (Table S4). As the seeding void is the sole area of contact between the cells and the matrix, we ignore the mechanical response of the mesh elements beyond the seeding void. The DNN is a fully connected feed-forward network with 4 hidden layers, each comprising 1,024 neurons with a ReLU activation function, and is designed to predict the Cartesian nodal displacement field of the seeding surface as outputs when given the Cartesian nodal force field exerted on the seeding surface as inputs. 6,000 samples of synthetic training data were generated for the training and validation process using the Abaqus finite element software, with the synthetic training data generation pipeline and DNN hyperparameters detailed in Supplementary Note 4-5. The displacement, stress and strain fields of the mesh nodes beyond the seeding surface are calculated as a post-processing step - the displacement field of the seeding surface, predicted from DNN-FE, is applied as a boundary condition in Abaqus, allowing for the displacement, and thus stress and strain, of the other mesh nodes to be computed. Post-processing is performed after the simulation has terminated to reduce computational overhead.

### Contact model

In our combined cell-matrix model, we focus exclusively on mechanical forces arising cell-cell and cell-matrix interactions, and we ignore other biomechanical and biochemical phenomena such as cell-matrix adhesion and matrix degradation. To translate these interactions into our computational framework, we utilise a linear force contact model that generates repulsive force between the cells and the ECM, scaling linearly with overlap distance to convert cell-ECM mechanical interactions to nodal forces *F*^*ECM*^ on the ECM. This contact model is described by 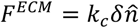, where *k*_*c*_ is the contact stiffness, *δ* is the overlap distance of the cell and matrix surface elements. We implement a viscous soft-relaxation method to maintain stable contact convergence between the proliferating cells and the ECM. The nodal position field of the ECM 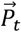 at time *t* is updated according to 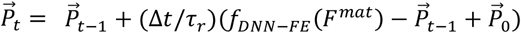. Here 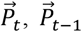, and 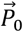 are the nodal position fields at time *t, t* − 1, and 0 respectively, Δ*t* is the simulation timestep and *τ*_*r*_ is the soft relaxation viscosity. The DNN-FE framework is denoted by *f*_*DNN*−*FE*_ (*F*^*mat*^).

### T47D spheroid culture

Human breast cancer T47D cells were expanded in media (Roswell Park Memorial Institute - RPMI) supplemented with 10% FBS, 100 U/mL penicillin and 100 mg/mL streptomycin. Cells were expanded at 37°C, in an incubator with 5% CO2 and culture media was replenished every 3 days. Gelatin-Transglutaminase hydrogels were prepared by mixing gelatin (type A, 175 Bloom, Sigma) in Roswell Park Memorial Institute (RPMI) 1640 culture medium containing 10% Fetal Bovine Serum (FBS) (Sigma) and 1% penicillin/streptomycin (Sigma) to a concentration of 9% w/v. The solution was heated at 80°C for 30 min, then sterile filtered using a 0.22 mm filter (Millipore). Microbial transglutaminase (mtgase) (Activa WM; containing 1% mtgase; Ajinomoto foods Europe S.A.S., France) was mixed with sterile phosphate buffered saline (PBS) before being sterile filtered and stored at −4°C until use. A suspension of Nano-hydroxyapatite (nHa) particles was created by mixing calcium chloride dihydrate solution and phosphate solution in a 1:1 v/v proportion. The hydrogels were formed by mixing gelatin, mtgase solution, and nHa solutions in a 1:1:1 (v/v:v) ratio. To provide hydrogels with varying stiffness, mtgase of different concentrations (0.3, 0.6, and 1% mtgase per gram of gelatin) were mixed with equal amounts of gelatin and nHa solutions. The final gelatin concentration for all the hydrogels was 3% w/v with 5.5% v/v nHa. The compressive moduli of the hydrogels were measured under unconfined compression in our previous study (Kumar et al., 2024). Cells were encapsulated within gelatin hydrogels of three different stiffnesses (0.58 kPa, 0.85 kPa,1.1 kPa) and hydrogel suspensions were pipetted into custom-made polydimethylsiloxane (PDMS) wells in a layer (1mm(height) x 4mm(breadth) x 13mm(length)) and cooled at 4°C for 8 min. The cell number was kept at 100,000 cells/hydrogel (2*x*10^6^ cells/mL of hydrogel). T47D-laden hydrogels were cultured in RPMI supplemented with 10% FBS, 100 U/mL penicillin and 100 mg/mL streptomycin. The media was changed every 2 days for the duration of the 3D culture period. All hydrogel groups were cultured for 12 days to facilitate spontaneous spheroid formation and growth. Fluorescent staining was performed on hydrogels to visualize the actin cytoskeleton and nuclei of the cells, and the formation of T47D spheroids. Hydrogels were fixed on days 3, 7, and 12 using 4% paraformaldehyde at 4°C overnight. Cells within these hydrogels were then permeabilized with 0.5% Triton X-in PBS for 10 min at 4°C under agitation. Samples were stained with phalloidin-FITC at 1.25 mg/mL (1:400) to stain the actin cytoskeleton and DAPI dilactate (1:2000) to identify cell nuclei, respectively. Z stack imaging was carried out using a Fluoview FV1000 confocal laser scanning microscope system (Olympus) at a magnification of 10x (air) and 20x (oil) with a step size of 5 mm. All stacks were obtained at the same intensity setting between groups. NIH ImageJ software was used to analyse stacks of images at maximum intensity projections. Spheroid diameters and cell numbers were measured from these maximum intensity projection images. Assuming the spheroid is perfectly spherical, average cell volume was estimated by dividing the spheroid volume by the population. Statistical analyses performed using GraphPad Prism software (version 8.4.3). Two-way ANOVA was used for analysis of variance with Bonferroni’s post-hoc tests to compare between different timepoints and experimental groups. Two replicates were analysed for each experimental group.

## Acknowledgements

The authors would like to acknowledge financial support from the Irish Research Council (GOIPG/2022/910) and the European Research Council (Starting Grant: 101116234 and Consolidator Grant: 863795). The work was also supported by funding from the University of Galway DISC PhD Scholarship. This publication has also emanated from research supported by the Irish Research Council Laureate Award Programme (MEMETic, IRCLA/2017/217), and Research Foundation Flanders (11D9923N). The authors would like to acknowledge the Centre for Microscopy and Imaging facilities at the University of Galway, including Prof Peter Owens and Dr Kerry Thompson.

## Data Availability Statement

All code and data will be made available on GitLab prior to peer-reviewed publication of the article.

## Author Contributions

I.S. and E.Mc. designed the theoretical models; I.S. and J.V. implemented the models and carried out the computations; V.K. and L.Mc. designed and conducted the experiments; I.S. designed and implemented the DNN-FE models; All authors (I.S., J.V, V.K, L.Mc., B.S., E.H., and E.Mc.) analysed and interpreted the data; I.S., J.V, B.S., E.H., and E.Mc wrote the initial manuscript. All authors (I.S., J.V, V.K, L.Mc., B.S., E.H., and E.Mc.) contributed to manuscript reviewing and editing.

