## Supplementary Information for "Stress-dependent growth of breast cancer models arises from a cellular volume checkpoint"

**Supplementary Note 1. Extended description of single cell cortex model.**

Water influx driven by hydrostatic and osmotic pressure gradients leads to an increase in cell volume, stretching the cell cortex. In this study, we treat the actomyosin cortex and membrane as a continuous mechanically unified structure, denoted as the cortex, and we neglect the possibility of blebbing and cortical detachment. The total cortical stress for a single spherical cell $i$ can be written as $\sigma_i = \sigma_{p,i} + \sigma_a$, where $\sigma_{p,i}$ is the passive stress arising from the deformation of the actin network and $\sigma_a$ is the active stress arising from myosin contractility. Existing cell models describe the cortex exhibiting both viscous and elastic properties, such that the passive stress may be expressed as $\sigma_{p,i} = k(A_i/A_o - 1)/2 + \eta(1/A_i)(dA_i/dt)$, where $k$ is the cortical stiffness, $A_i$ and $A_o$ are the current and reference cell surface areas respectively, and $\eta$ is the cortex viscosity. Experimental evidence suggests the cortex viscosity to lie in the range of $10^2$-$10^3$ Pa/s (Evans & Yeung, 1989; Koay et al., 2003) with the cell radius shrinking by 10% over several minutes under load (Stewart et al., 2011). Arising from the insignificant viscous contribution $\eta(2/r)(dr/dt) \approx 0.1 - 1$ Pa to the passive stress, we neglect the contribution of the viscous component, with a view that k denotes the long-term cortical stiffness. Therefore, the total stress reduces to $\sigma_i = k(r_i^2/r_0^2 - 1)/2 + \sigma_{a,}$. The cell experiences mechanical loading arising from a uniform external fluid pressure $P^{ext}$, with the pressure difference across the cell membrane denoted as $\Delta P_i = P_i - P^{ext}$. From Laplace's law, the cortical stress can also be written as $\sigma_i = \Delta P_i r_i/2h$, where $h$ is the cortical thickness. In addition, mechanical interactions with surrounding cells and the extracellular matrix (ECM) generate contact stress $\sigma_{ext}$. Thus, the constitutive law and mechanical balance forms of the cortical stress are equivalent:

$$\sigma_i = k/2(r_i^2/r_0^2 - 1) + \sigma_a = (\Delta P_i - \sigma_{ext})r_i/2h \,. \qquad (S1)$$

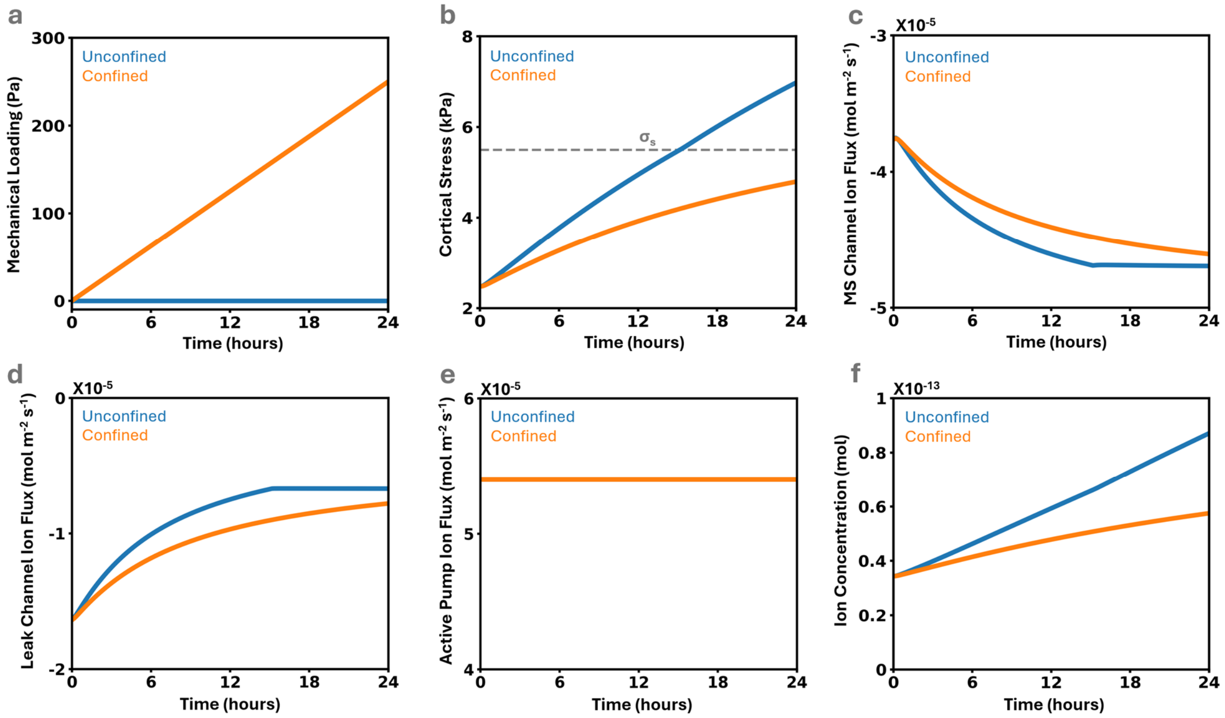

***Figure S1: Additional figures for single cell analysis under confinement.*** *(a) Form of confinement stress applied. (b) Differences in cortical stress under confinement. Differences in ion flux through (c) mechanosensitive ion channels, (d) leak channels, and (e) active ion pumps under confinement. (f) Difference in permeable ion concentration under confinement.*

**Supplementary Note 2. Numerical implementation of deformable cell model.**

In this section we provide the implementation of the discretized active foam model previously described by (Cuvelier et al., 2023; Ongenae et al., 2024; Vangheel et al., 2025). In this model, cells are represented as pressurized bubbles with a viscous cortex under tension and adhesive bonds allowing cell-cell and cell-substrate contact. The cell actomyosin complex is represented by a triangulated surface mesh where the vertex positions $\mathbf{x}_i$ act as the relevant degrees of freedom. Associated parameters are summarised in Supplementary Note 3.

**Cortex tension.** Tension in the cortex arises from active actomyosin contractility $\gamma_\alpha$ and a strain dependent tensile modulus as $\gamma_{cm} = \gamma_\alpha + K_\gamma \epsilon_S$, with $\gamma_a$the active base tension, $K_\gamma$the area modulus, and $\epsilon_S = (S_i - S_{0,i})/S_{0,i}$ the area strain of the cell with instantaneous area $S_i$ and reference area $S_{0,i}$. At contacting interfaces, cortex tension decreases with a reduction in actomyosin activity (Maître et al., 2012) represented as an effective adhesive tension $\omega$ as $\gamma_I = \gamma_{cm} - \omega$. The resulting surface tension in a triangle $A$ is given by

$$\gamma_c = \begin{cases} \gamma_I & \text{for contacting surfaces} \\ \gamma_{cm} & \text{for free surfaces.} \end{cases} \tag{S2}$$

Within this framework, cortical tension can be equivalently expressed as a surface (cortical) stress σ, given by

$$\sigma = \frac{\gamma_c}{h}, \tag{S3}$$

where $h$ is the cortical thickness. Assuming tension balance, the contact angle between two adhering cells can be computed as $\cos\theta = 1 - \omega/\gamma_{cm}$. We assume this contact angle is constant during cell growth, so we scale adhesive tension $\omega$ with cell medium tension $\omega_t = \omega/\gamma_{cm}$. The force on each node $i$ of a triangle consisting out of nodes $A = (i, j, k)$ is then given by (Fedosov et al., 2010)

$$\mathbf{F}_i^{act} = \frac{\gamma_c}{2}\left(\mathbf{x}_k - \mathbf{x}_j\right) \times \hat{\mathbf{n}}_A. \tag{S4}$$

**Cortex viscosity.** Actomyosin remodelling is described by an effective viscosity $\eta_c$. We distinguish between in-plane and out-of-plane bending viscosity. The viscous damping force between two nodes $i, j$ with velocity $\boldsymbol{v}$ is given by

$$\mathbf{F}_\mathbf{i}^{\text{visc}} = \frac{\eta_c t_c}{\sqrt{3}}\left(\left(\hat{\mathbf{n}}_{ij} \cdot (\mathbf{v}_j - \mathbf{v}_i)\right)\hat{\mathbf{n}}_{ij} + \left(\hat{\mathbf{t}}_{ij} \cdot (\mathbf{v}_j - \mathbf{v}_i)\right)\hat{\mathbf{t}}_{ij}\right) + \left(\hat{\mathbf{b}}_{ij} \cdot (\mathbf{v}_j - \mathbf{v}_i)\right)\hat{\mathbf{b}}_{ij}, \tag{S5}$$

where the direction of viscous forces are given by the unit vectors

$$\hat{\mathbf{n}}_\mathbf{ij} = \frac{\mathbf{x_j} - \mathbf{x_i}}{\| \mathbf{x_j} - \mathbf{x_i} \|}, \tag{S6}$$

$$\hat{\mathbf{b}}_{ij} = \frac{\hat{\mathbf{n}}_A + \hat{\mathbf{n}}_B}{\|\hat{\mathbf{n}}_A + \hat{\mathbf{n}}_B\|}, \tag{S7}$$

$$\hat{\mathbf{t}}_\mathbf{ij} = \hat{\mathbf{b}}_{ij} \times \hat{\mathbf{n}}_{ij}, \tag{S8}$$

with $\hat{\mathbf{n}}_\mathbf{A}$, and $\hat{\mathbf{n}}_\mathbf{B}$, the normals of the triangles making up the segment between nodes $i$ and $j$. Note that we do not distinguish between bulk, extensional ($\eta_e$) or shear viscosity ($\eta_c$), effectively setting the Trouton's ratio to $\eta_e/\eta_c = 1$, whereas a Newtonian fluid would have a value equal 3.

**Cytokinesis.** Cell division and cytokinesis is modelled by applying an elastic ring that shrinks during the cytokinesis process, and eventually splits the mother cell into two cells. We first define the direction of cell division $\mathbf{p}$, which separates the nodes $i$ of the mother cell into two daughterhalves ($D_1$ and $D_2$) as

$$\begin{aligned} \mathbf{p} \cdot (\mathbf{x}_i - \mathbf{x}_c) \geq 0 \rightarrow i \in D_1, \\ \mathbf{p} \cdot (\mathbf{x}_i - \mathbf{x}_c) < 0 \rightarrow i \in D_2, \end{aligned} \quad (S9)$$

where $\mathbf{x}_c$ is the cell centre. The nodes that create the boundary between the two halves eventually make up the constriction ring. The elastic force on connecting nodes $i, j$ of this ring is given by

$$\mathbf{F}_i^{div} = k_r \left[ \frac{l_{s,ij}}{l_{0,ij}} - \left(1 - \frac{t}{\tau_d}\right) \right] \frac{\mathbf{x_j} - \mathbf{x_i}}{\parallel \mathbf{x_j} - \mathbf{x_i} \parallel}, \quad (S10)$$

where $k_r$ is the ring relative stiffness, $l_s$ is the instantaneous length of the segment between two nodes, $l_0$ the initial segment length before cytokinesis, and $\tau_d$ is the duration of cytokinesis. Finally, when the constriction ring reaches the abscission radius $r_a$, the cell is split into two daughter cells. Abscission is performed by adding two new nodes – one for each daughter – at the centre of the mother cell, and connecting these nodes with the nodes from the constriction ring obtaining two daughter cells.

**Hydrostatic pressure.** We assume the cell cytoplasm to be incompressible, with a volume governed by our hydromechanical model (eqn 1). Therefore, we implement active volume control for each cell by a cytoplasmic pressure $P_b$ using a PI volume controller

$$P_b^{cyt} = -K_p \epsilon_V(t) - K_i \int_0^t \epsilon_V(t) \mathrm{dt}, \quad (S11)$$

where $\epsilon_V(t) = (V'(t) - V(t))/V(t)$, with $V'(t)$ the instantaneous volume and $V(t)$ is the target volume, $K_p$ the proportional and $K_i$ integrative term. Furthermore, during cytokinesis the daughter halves are stabilized by a secondary pressure

$$P_b^{div} = K_d \left(1 - \frac{V_d}{V_d^*}\right), \quad (S12)$$

with equilibrium volume $V_d^*$ and instantaneous volume $V_d$. The total pressure from volume controll acting on a node is then given by $P_b(t) = P_b^{cyt} + P_b^{div}$. The force on a triangle $A$ is given by

$$\mathbf{F}_A^b = \mathbf{S}_A P_b(t), \quad (S13)$$

with $\mathbf{S}_A$ the directed area of the triangle, that is simply distributed to the nodes $i$ as

$$\mathbf{F}_i^b = \frac{\mathbf{F}_A^b}{3}. \quad (S14)$$

**Contact pressure.** We assume uniform adhesion over interacting surfaces. The contact pressure $P_{AB}^{c}$ includes repulsive and adhesive interactions as

$$P_{AB}^{c}(\mathbf{x}) = -k_c\delta(\mathbf{x}) + P^0, \tag{S15}$$

where $P^0 = \omega/h_0$, with $h_0$ the effective range of adhesion that is accomplished by translating the nodes over the distance $h_0/2$ in the direction of the node normals. To ensure that at equilibrium the work of adhesion is recovered, the stiffness is set as $k_c = P^0/h_0$. The contact overlap at point $\mathbf{x}$ on a triangle is computed with the translated triangles as

$$\delta(\mathbf{x}) = \max(0; 2(\mathbf{x} - \mathbf{s}_{AB}) \cdot (\hat{\mathbf{n}}_{AB} \times \mathbf{l}_{AB}) \tan(\alpha)), \tag{S16}$$

with $\mathbf{l}_{AB}$ the intersection line between two triangles, $\mathbf{s}_{AB}$ an arbitrary point on the intersection line, $\hat{\mathbf{n}}_{AB}$ the normal of the common contact plane defined as

$$\hat{\mathbf{n}}_{AB} = \frac{\hat{\mathbf{n}}_A - \hat{\mathbf{n}}_B}{\| \hat{\mathbf{n}}_A - \hat{\mathbf{n}}_B \|}, \tag{S17}$$

and $\alpha$ the angle between the triangle and the contact plane $A \cap B$. The contact force and moment is obtained by integrating the total contact pressure over the oriented contact area $\mathbf{S}_{AB}$ on the common contact plane

$$\mathbf{F}_{AB}^{c} = \int_{\mathbf{x}\in A\cap B} P_{AB}^{c}(\mathbf{x}) d\mathbf{S}_{AB}, \tag{S18}$$

$$\mathbf{M}_{AB}^{c} = \int_{\mathbf{x}\in A\cap B} P_{AB}^{c}(\mathbf{x}) d\mathbf{S}_{AB} \times (\mathbf{x} - \mathbf{x}_{AB}), \tag{S19}$$

with $\mathbf{x}_{AB}$ the contact point defined as the geometric center of the intersection polygon. This force is distributed to the nodes $i$ ensuring conservation of momentum and force, assuming the forces are directed in $\hat{\mathbf{n}}_{AB}$. This results in a system of linear equations per contact pair

$$\mathbf{M}_{AB}^{c} = \sum_{i\in A} (\mathbf{x} - \mathbf{x}_{AB}) \times \mathbf{F}_{i,AB}^{c}, \tag{S20}$$

$$\mathbf{F}_{AB}^{c} = \sum_{i\in A} \mathbf{F}_{i,AB}^{c}. \tag{S21}$$

The solution for this system is presented and discussed in the work of Smeets et al., (2015).

**Wet contact friction.** A viscous contact force is included to account for drag between contacting surfaces. The contact drag force acting on node $i$ of triangle $A$ of the contact pair $(AB)$ is computed as

$$\mathbf{F}_{AB,i}^{\text{fric}} = \Gamma_{AB}^{\text{fric}} \cdot \sum_{k\in B} w_{ik}^{AB} (\mathbf{v}_k - \mathbf{v}_i), \tag{S22}$$

determined by a friction tensor $\Gamma_{AB}^{\text{fric}}$ and weights $w_{ik}^{AB}$ per node $k$ of the $B$ triangle. $w_{ik}^{AB}$ are assumed to scale with the relative contribution of the nodal contact forces to the overall contact force thus

$$w_{ik}^{AB} = \frac{\left(\mathbf{F}_{AB,i}^{c} + \mathbf{F}_{AB,k}^{c}\right) \cdot \hat{\mathbf{n}}_{AB}}{6\sum_{\forall k \in B} \mathbf{F}_{AB,k}^{c} \cdot \hat{\mathbf{n}}_{AB}} \qquad (S23)$$

is used as an approximation. $\Gamma_{AB}^{\text{fric}}$ for a given contact area $S_{AB}$ is estimated as

$$\Gamma_{AB}^{fric} = S_{AB}[\xi^{\perp}\hat{\mathbf{n}}_{AB} \otimes \hat{\mathbf{n}}_{AB} + \xi^{\parallel}(-\hat{\mathbf{n}}_{AB} \otimes \hat{\mathbf{n}}_{AB})], \qquad (S24)$$

with normal and tangential friction coefficients $\xi^{\perp}$ and $\xi^{\parallel}$ respectively.

**Medium damping.** A general drag force on node $i$ from liquid drag between cell and medium with viscosity $\eta_f$ is given by

$$\mathbf{F}_i^{\text{drag}} = -\Gamma_i^{\text{f}} \cdot \mathbf{v}_i, \qquad (\text{S25})$$

where we approximate the drag force from Stokes' Law for spherical particles as

$$\Gamma_i^{\text{f}} = 6\pi R_0 \eta_f \frac{S_i}{4\pi R_0^2} = \frac{3\eta_f}{2R_0} S_i. \qquad (\text{S26})$$

$F_i^{\text{drag}}$ is small compared to other dissipative forces but provides stability of the numerical integration scheme as it ensures the resistance matrix is positive definite.

**Equation of motion.** For the overdamped environment, inertial forces can be neglected and the force balance for node $i$ is given by

$$\mathbf{F}_i^{\text{act}} + \mathbf{F}_i^{\text{b}} + \sum_j \mathbf{F}_{ij}^{div} + \sum_{(AB):i\in A} \mathbf{F}_{i,AB}^{c} = -\mathbf{F}_i^{\text{drag}} - \sum_j \mathbf{F}_{ij}^{\text{visc}} - \sum_{(AB):i\in A} \mathbf{F}_{i,AB}^{\text{fric}}, \qquad (\text{S27})$$

Where the left-hand forces include force from cortex tension, bulk pressure, constriction ring summed over all connecting nodes $ij$, and contact forces summed over all contacting triangles AB. Right-hand forces are viscous forces including medium drag, viscous cortex damping summed over connecting nodes $ij$, and friction forces summed over contacting triangles $AB$. These viscous forces linearly depend on relative velocities of the nodes. For the total system of $N$ nodes, this can be written as

$$\mathbf{F} = \mathbf{\Lambda}\,\mathbf{v}, \qquad (\text{S28})$$

with $\mathbf{F}$ a $3N \times 1$ column matrix that represents all left-hand forces per node, $\mathbf{\Lambda}$ a $3N \times 3N$ sparse symmetric positive definite friction matrix that contains off-diagonal terms that capture cortical viscosity and contact friction, and $\mathbf{v}$ a $3N \times 1$ column matrix representing the velocities of $N$ nodes. The congujate gradient method (CGM) is then used to solve for the node velocities every timestep. Node positions $\mathbf{x}_i$ are updated every timestep using a semi-implicit Euler integration scheme

$$\mathbf{x}_i(t + \Delta t) = \mathbf{x}_i(t) + \Delta t\,\mathbf{v}_i(t + \Delta t), \qquad (\text{S29})$$

where $\mathbf{v}_i(t + \Delta t)$ are the projected new velocities obtained by solving eqn S28 at time $t$ via the CGM.

**Remeshing.** As we model the cortex as a viscous fluid under surface tension, the discrete nodes tend to flow over the surface. To ensure the mesh quality, we remesh the triangulated surface. There are 4 remeshing stages based on (Brakke, 1992): split triangles, flip edges, split edges, and delete edges. In the first stage, triangles that are too big ($S_A > S_{min}$) are split at the geometric center in 3 separate triangles. In the second stage, edges between triangles are flipped if the angle of two vertices opposing the edge add up to more than $\pi$. For a curved surface, this operation can result in area, volume and shape changes. To limit this effect, flipping is only done for edges between neighbouring triangles that make an angle $\theta < \pi/3$. Otherwise, the edge is split in the center where both triangles are subdivided into two triangles. In the last stage, edges between triangles that are too small ($S_A < S_{max}$) are deleted, removing both triangles. This is repeated every $2\eta_c t_c/\sqrt{3}\gamma\Delta t$ timesteps. With remeshing, we assume that the nodes do not carry local information, but are merely considered as discretization points to represent a continuous surface.

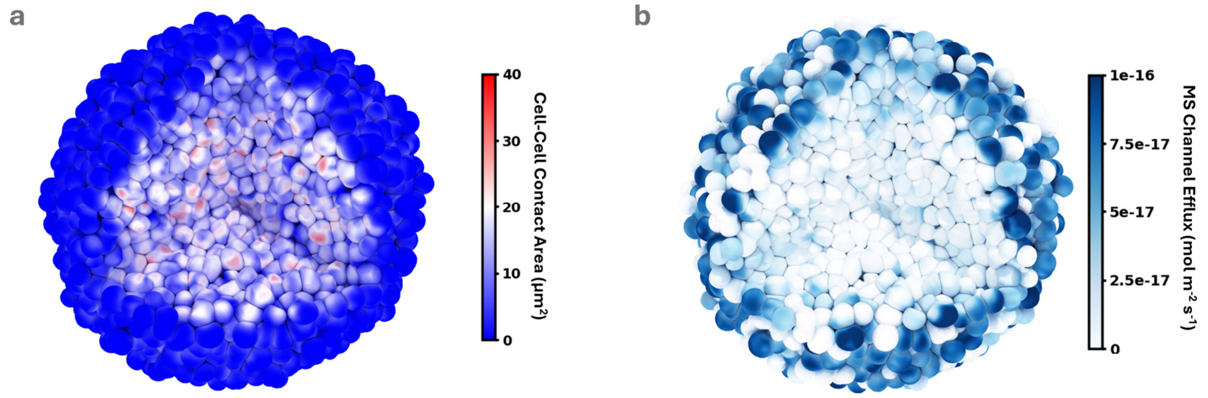


***Figure S2: Additional figures for single cell analysis under confinement. (a)*** *A volume cutout exposes the* ***(a)*** *increase in cell-cell contact area at the core, and the* ***(b)*** *corresponding reduction in ion flux through the MS channels.*

**Supplementary Note 3. Motivations for model parameters.**

Here we first outline modelling parameters and motivation for single cell analyses. The membrane solvent permeability $P_s$ has been reported to lie between $10^{-4}$ – $10^{-5}$ m/s for a range of cell types. Considering the molar volume of water $\bar{V}_W$ = 18 mL/mol and following from Jiang & Sun, (2013), the water permeability of the membrane is expressed as $L_{p,m} = P_s\bar{V}_W/RT$, where $R$ is the gas constant and $T$ is the absolute temperature. We obtain an upper value of $L_{p,m}$ = 7x$10^{-12}$ m.$s^{-1}$$Pa^{-1}$. Based on experimental evidence (Clark et al., 2013; Laplaud et al., 2021), we define the thickness and stiffness of the cell cortex as 250 nm and 10 kPa respectively. In line with Jiang & Sun, (2013), we assume $\sigma_a$ = 100 Pa. Following from McEvoy et al., (2020), the external osmotic pressure $\Pi^{ext}$ = 0.67 MPa and the critical osmotic pressure for active pumping $\Delta\Pi_c$ = 40 GPa with $\zeta = 2.25x10^{-9}$ mol.$m^{-2}$$s^{-1}$$Pa^{-1}$. As ion flux across the mechanosensitive and leak channels has been reported to lie in the range of $10^{-7}$ – $10^{-6}$ mol.$m^{-2}$$s^{-1}$ (Grosell, 2006; Larsen et al., 2007), we choose the permeability of the mechanosensitive ion channels and leak channels as $\beta$ = 1.05x$10^{-8}$ mol.$m^{-2}$$s^{-1}$$Pa^{-1}$ and $\varpi_l$ = 1.5x$10^{-9}$ mol.$m^{-2}$$s^{-1}$$Pa^{-2}$ respectively. Based on experimental data measuring the growth curve of single HeLa cells up to the point of mitosis (Cadart et al., 2022),we approximate a value of $V_{crit}$ as 2700 μ$m^3$. Single-cell hydromechanical model parameters for all simulations are summarised in Table S1.

| Parameter | Description | Value |
|---|---|---|
| $r_0$ | reference cell radius ($m$) | 5.5x$10^{-6}$ |
| $k$ | cortex layer stiffness ($Pa$) | 1x$10^{4}$ |
| $\sigma_a$ | active cortical stress ($Pa$) | 100 |
| $h$ | cortical layer thickness ($m$) | 2.5x$10^{-7}$ |
| $\beta$ | mechanosensitive channel ion permeability ($mol.m^{-2}s^{-1}Pa^{-1}$) | 1.05x$10^{-8}$ |
| $\sigma_c$ | mechanosensitive channel threshold stress ($Pa$) | 1000 |
| $\sigma_s$ | mechanosensitive channel saturation stress ($Pa$) | 5500 |
| $\varpi_l$ | leak channel ion permeability ($mol.m^{-2}s^{-1}Pa^{-1}$) | 1.05x$10^{-9}$ |
| $\zeta$ | active pumping rate constant ($mol.m^{-2}s^{-1}Pa^{-1}$) | 2.25x$10^{-17}$ |
| $\Delta\Pi_c$ | active pumping critical osmotic pressure difference ($Pa$) | 4x$10^{10}$ |
| $\Pi^{ext}$ | mechanosensitive channel threshold stress ($Pa$) | 1.05x$10^{-9}$ |
| $L_{p,m}$ | cell membrane water permeability ($m.s^{-1}Pa^{-1}$) | 0.7x$10^{-11}$ |

***Table S1:*** *Parameters for single-cell hydromechanical cell model.*

Relevant parameters from single cell analyses are maintained for multicellular simulations unless otherwise stated. For T47D analyses, we approximate a value of $V_{crit}$ as 4300 μ$m^3$, allowing for the distribution of average cell volumes to align with our experimental data. The proliferative capability parameter $\lambda = 1.25\text{x}10^{-5}$ , synthesis rate $\alpha_x = 5.83x10^{-21}$ mol/s, and biomolecule saturation constant $N_{xs} = 1.5\text{x}10^{-15}$ mol have been tuned to allow predictions to align with experimental T47D data with a stochastic mitosis parameter $z = 20$. Numerical parameters for the Mpacts framework are constrained to previously reported ranges, as follows. Tension in the cortex arises from actin tension with $\omega_t = 0.25$ and $K_\gamma = 1.8$ nN/μm, with remodelling described through viscous dissipation with $\eta_c = 3x10^4\ Pa.s$. Cytokinesis has a characteristic timescale $\tau_d = 60\ s$ and an associated ring stiffness $k_r = 2.21\ x\ 10^{-7}\ kg.m/s^2$ and abscission radius $r_a = 9.1x10^{-7}\ m$. Target volume is achieved through a proportional integrative controller (eqn S11) with control terms $K_p = 1x10^4$ Pa and $K_i = 0.25\ s$. The stiffness of a nodal contact interaction is $k_c = 0.05\ kg/s^2$ with a relaxation timescale $\tau_r = 60\ s$ and an effective range $h_0 = 2x10^{-7}\ m$, and there are additional normal and tangential frictional interactions with coefficients $\xi^{\perp} = \xi^{\parallel} = 1.33x10^9 kg/(m^2.s)$, respectively. The surrounding medium is assumed to have a viscosity $\eta_f = 1x10^3$ Pa.s. For the surface mesh, the max and min triangle areas are $S_{max} = 1.81x10^{11}\ m^2$ and $S_{min} = 2.01x10^{12}\ m^2$, respectively. These multi-cellular Mpacts framework parameters are summarised in Table S2. For HeLa and 4T1 cells, the growth parameters are modified such that $V_{crit}^{HeLa} = 2700\ \mu m^3$; $V_{crit}^{4T1} = 3300\ \mu m^3$ , $\lambda^{HeLa} = 0.5\text{x}10^{-4}$; $\lambda^{4T1} = 1\text{x}10^{-4}$, $\alpha_x^{HeLa} = 4.58x10^{-21}$ mol/s; $\alpha_x^{4T1} = 6.67x10^{-21}$ mol/s, and $N_{xs}^{HeLa} = 5\text{x}10^{-16}$ mol; $N_{xs}^{4T1} = 9\text{x}10^{-16}$ mol.

| Parameter | Description | Value |
|---|---|---|
| $\alpha_x$ | biomolecule synthesis rate ($mol\ s^{-1}$) | $5.83\text{x}10^{-21}$ |
| $N_{xs}$ | biomolecule saturation constant ($mol$) | $1.5\text{x}10^{-15}$ |
| $V_{crit}$ | critical mitotic volume ($\mu m^3$) | 4300 |
| $\lambda$ | proliferative capacity ($-$) | $1.25\text{x}10^{-5}$ |
| $\varrho$ | mitosis stochasticity ($-$) | 20 |
| $\gamma_\alpha$ | cortex active tension ($nN/\mu m$) | 0.06 |
| $K_\gamma$ | tension modulus ($nN/\mu m$) | 1.8 |
| $S_0$ | cell reference area ($\mu m^2$) | $4\pi r_0^2$ |
| $\omega_t$ | relative adhesive tension ($-$) | 0.25 |
| $\omega_{t,div}$ | relative adhesive tension dividing cells ($-$) | 0.05 |
| $\eta_c$ | cortex viscosity ($Pa.s$) | $3\text{x}10^4$ |
| $k_r$ | ring relative stiffness ($kg.m.s^{-2}$) | $2.21\text{x}10^{-7}$ |
| $\tau_d$ | cytokinesis duration ($s$) | 60 |
| $r_a$ | abscission radius ($m$) | $9.1\text{x}10^{-7}$ |
| $K_p$ | proportional volume control term ($Pa$) | $1\text{x}10^4$ |
| $K_i$ | integrative volume control term ($s$) | 0.25 |
| $h_0$ | effective adhesion range ($m$) | $2\text{x}10^{-7}$ |
| $\xi^{\perp}$ | normal friction coefficient ($kg.m^{-2}.s^{-1}$) | $1.33\text{x}10^9$ |
| $\xi^{\parallel}$ | tangential friction coefficient ($kg.m^{-2}.s^{-1}$) | $1.33\text{x}10^9$ |
| $\eta_f$ | medium viscosity ($Pa.s$) | $1\text{x}10^3$ |
| $S_{\min}$ | minimum triangle area ($m^2$) | $2.01\text{x}10^{-12}$ |

***Table S2:*** *Parameters for Mpacts multi-cellular framework.*

**Supplementary Note 4. Generation of synthetic training data.**

Arising from the use of deep learning to model the mechanical response of the matrix, the diversity of the dataset is of crucial importance. Diverse datasets allow for improved generalisation of the model and reduced bias (Alruwaili & Alsalim, 2024; Yu et al., 2022). Each training sample generates a random nodal force field applied on the seeding surface (Alg. 1). The nodal forces are randomly applied in the radial direction away from the centre of the seeding void, with $D_f = [[-0.05, 0.05], [-0.05, 0.05], [1, 1]]$ to apply 5% variance in the orthogonal directions to account for shear forces applied by cells onto the ECM. To ensure the training data covers a range of loading scenarios and edge cases, the training samples are divided into subsets, where each subset provides a number of training samples for a particular loading case (Table S3). 6,000 synthetic training samples are created using the Abaqus finite element software (version 2021). As the seeding surface is the sole point of contact between the proliferating cells and the ECM, we only consider the nodal force and displacement fields of the seeding surface in the DNN-FE framework, minimising the number of parameters in the deep learning model. The network inputs and outputs are normalised using min-max normalisation to improve model convergence.

**Algorithm 1** Generation of a random nodal force field for each training sample

**procedure** GENERATERANDOMNODALFORCEFIELD($P, A, \mathbf{o}, D_f, F_{lim}, P_f$)

$P \in \mathbb{R}^{n\times 3}$ — Seeding surface node positions
$A \in \mathbb{R}^{n}$ — Seeding surface node areas
$\mathbf{o} \in \mathbb{R}^{3}$ — Centre of seeding surface
$D_f \in \mathbb{R}^{3\times 2}$ — Bounds of force distribution per component
$F_{lim} \in \mathbb{R}^{2}$ — Bounds of force magnitude
$P_f \in \mathbb{R}$ — Force application probability

$A_{max} \leftarrow \max(A)$
$\mathbf{F} \in \mathbb{R}^{n\times 3} \leftarrow 0$ ▷ Nodal force field initialised as zero forces
**for** $i \in \{1,\ldots,n\}$ **do**
$\mathbf{p}_i \leftarrow \mathbf{P}[i]$ ▷ Node position
$a_i \leftarrow A[i]$ ▷ Node area
$\mathbf{v} \leftarrow \left[x \sim \mathscr{U}\left((D_f)_{1,1},(D_f)_{1,2}\right),\ y \sim \mathscr{U}\left((D_f)_{2,1},(D_f)_{2,2}\right),\ z \sim \mathscr{U}\left((D_f)_{3,1},(D_f)_{3,2}\right)\right]$
$\mathbf{d}_f \leftarrow \frac{\mathbf{v}}{\|\mathbf{v}\|}$ ▷ Force distribution vector
$\mathbf{f} \leftarrow$ COMPUTENODALFORCEVECTOR$(\mathbf{o}, \mathbf{p}_i, \mathbf{d}_f, F_{lim}) \cdot \frac{a_i}{A_{max}}$ ▷ Nodal force
$\mathbf{F}[i] \leftarrow \mathbf{f}_i$ **if** $r \sim \mathscr{U}(0,1) < P_f$ ▷ Nodal forces have a probability of being activated
**end for**
**return F**
**end procedure**

**procedure** COMPUTENODALFORCEVECTOR($\mathbf{p}_{\text{start}}, \mathbf{p}_{\text{end}}, \mathbf{d}_f, F_{lim}$)

$\mathbf{p}_{\text{start}} \in \mathbb{R}^3$ — Start position of force vector
$\mathbf{p}_{\text{end}} \in \mathbb{R}^3$ — End position of force vector
$\mathbf{d}_f \in \mathbb{R}^3$ — Force distribution per component
$F_{\text{lim}} \in \mathbb{R}^2$ — Bounds of force magnitude

$\mathbf{u} \leftarrow \frac{\mathbf{p}_{\text{end}} - \mathbf{p}_{\text{start}}}{\|\mathbf{p}_{\text{end}} - \mathbf{p}_{\text{start}}\|}$
**if** $\mathbf{u} \approx [1,0,0]$ **then** ▷ Create a non-parallel reference vector
$\mathbf{a} \leftarrow [0,1,0]$
**else if** $\mathbf{u} \approx [-1,0,0]$ **then**
$\mathbf{a} \leftarrow [0,-1,0]$
**else**
$\mathbf{a} \leftarrow [1,0,0]$
**end if**
$\mathbf{v}_1 \leftarrow \frac{\mathbf{u}\times\mathbf{a}}{\|\mathbf{u}\times\mathbf{a}\|}$ ▷ Orthogonal to **u**
$\mathbf{v}_2 \leftarrow \frac{\mathbf{u}\times\mathbf{v}_1}{\|\mathbf{u}\times\mathbf{v}_1\|}$ ▷ Also orthogonal to **u** and $\mathbf{v}_1$
$\mathbf{f} \leftarrow \frac{d_{f1}\mathbf{v}_1 + d_{f2}\mathbf{v}_2 + d_{f3}\mathbf{u}}{\|d_{f1}\mathbf{v}_1 + d_{f2}\mathbf{v}_2 + d_{f3}\mathbf{u}\|} \cdot \mathscr{U}\left(F_{lim1}, F_{lim2}\right)$
**return f**
**end procedure**

| Sample Type | $F_{lim}$ | $P_f$ | Number of samples |
|---|---|---|---|
| 1 | [0, 0.5x10$^{-7}$] | 0.05 | 500 |
| 2 | [0, 0.5x10$^{-7}$] | 0.2 | 500 |
| 3 | [0, 0.5x10$^{-7}$] | 0.4 | 500 |
| 4 | [0, 0.5x10$^{-7}$] | 0.6 | 500 |
| 5 | [0, 0.5x10$^{-7}$] | 0.8 | 500 |
| 6 | [0, 0.5x10$^{-7}$] | 0.95 | 500 |
| 7 | [0.5x10$^{-7}$, 1x10$^{-7}$] | 0.5 | 750 |
| 8 | [0.5x10$^{-7}$, 1x10$^{-7}$] | 0.6 | 750 |
| 9 | [0.5x10$^{-7}$, 1x10$^{-7}$] | 0.8 | 750 |
| 10 | [0.5x10$^{-7}$, 1x10$^{-7}$] | 0.95 | 750 |
| **Total** | | | **6000** |

***Table S3:*** *Training sample subsets and parameters.*

| Young's Modulus | Bulk Modulus | Shear Modulus |
|---|---|---|
| 0.31 $kPa$ | 120 $Pa$ | 250 $Pa$ |
| 0.58 $kPa$ | 220 $Pa$ | 480 $Pa$ |
| 0.85 $kPa$ | 320 $Pa$ | 710 $Pa$ |
| 1.1 $kPa$ | 420 $Pa$ | 910 $Pa$ |
| 1.37 $kPa$ | 520 $Pa$ | 1110 $Pa$ |

***Table S4:*** *Material properties of hydrogels.*

**Supplementary Note 5. DNN model training and validation.**

The deep neural network consists of hour hidden layers, each comprising 1,024 fully connected neurons with the ReLU activation function. An activation function is omitted in the output layer as the network outputs are normalised between 0 and 1. For model training we utilise the Huber loss function - a hybrid of mean squared error and mean absolute error, to decrease model sensitivity to outliers. The loss function quantifies the error between the ground truth $y$ and the predicted outputs $f(x)$, guiding the optimisation process by minimising this error during training to improve model accuracy.

$$L_\gamma(a) = \begin{cases} \frac{1}{2}a^2 & \text{if } |a| \leq z \\ z\left(|a| - \frac{1}{2}z\right) & \text{if } |a| > z \end{cases} \tag{S30}$$

where $a = z - f(x)$ and $z$ is the threshold parameter to select between mean squared error and mean absolute error. A learning rate of 2.5e-7 and mini-batch gradient descent with a batch size of 32 samples is used for model training. The Adam optimizer used to efficiently update weights during training (Kingma & Ba, 2014).

**Supplementary Note 6. Extended data on T47D simulations.**

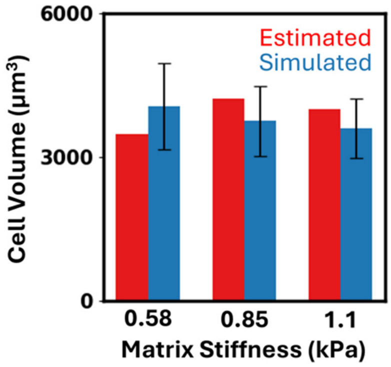


***Figure S3: T47D cell volumes.*** *Simulated T47D cell volumes align with estimations from experimental results. Experimental cell volumes estimated from mean spheroid volume / mean cells per spheroid. Error bars show mean ± SD.*

**Supplementary Note 7. Experimental validation of stress-dependent 4T1 spheroid growth.**

To further assess the predictive capabilities of our computational framework across a range of cell lines, we validate predictions against experimental growth of 4T1 murine mammary cells in gelatin hydrogels with varying Young's Moduli of 0.58 kPa, 0.85 kPa and 1.1 kPa (Kumar et al. 2024). As with the T47D validation, our framework predicts that an increase in matrix stiffness leads to an inhibition in cell proliferation and a subsequent reduction in spheroid size. Spheroid size is predicted to decrease by 30% as matrix stiffness increases from 0.58 kPa to 1.1 kPa, in close agreement with experimental data. Reflecting the trends observed in the T47D validation, the peak radial stress was similar in all three cases (~300 Pa, Fig. S4a). Predictions from our model are also consistent with experimentally-recorded spheroid diameter at all three timepoints (Fig. S4b), spheroid population (Fig. S4c) and experimentally-calculated average cell volume (Fig. S4d). Mean cytosolic hydrostatic pressure is predicted to increase with increasing hydrogel stiffness (Fig. S4e). This, in turn, reduces the rate of cell growth and restricts cells from surpassing the mitotic checkpoint, limiting spheroid expansion.

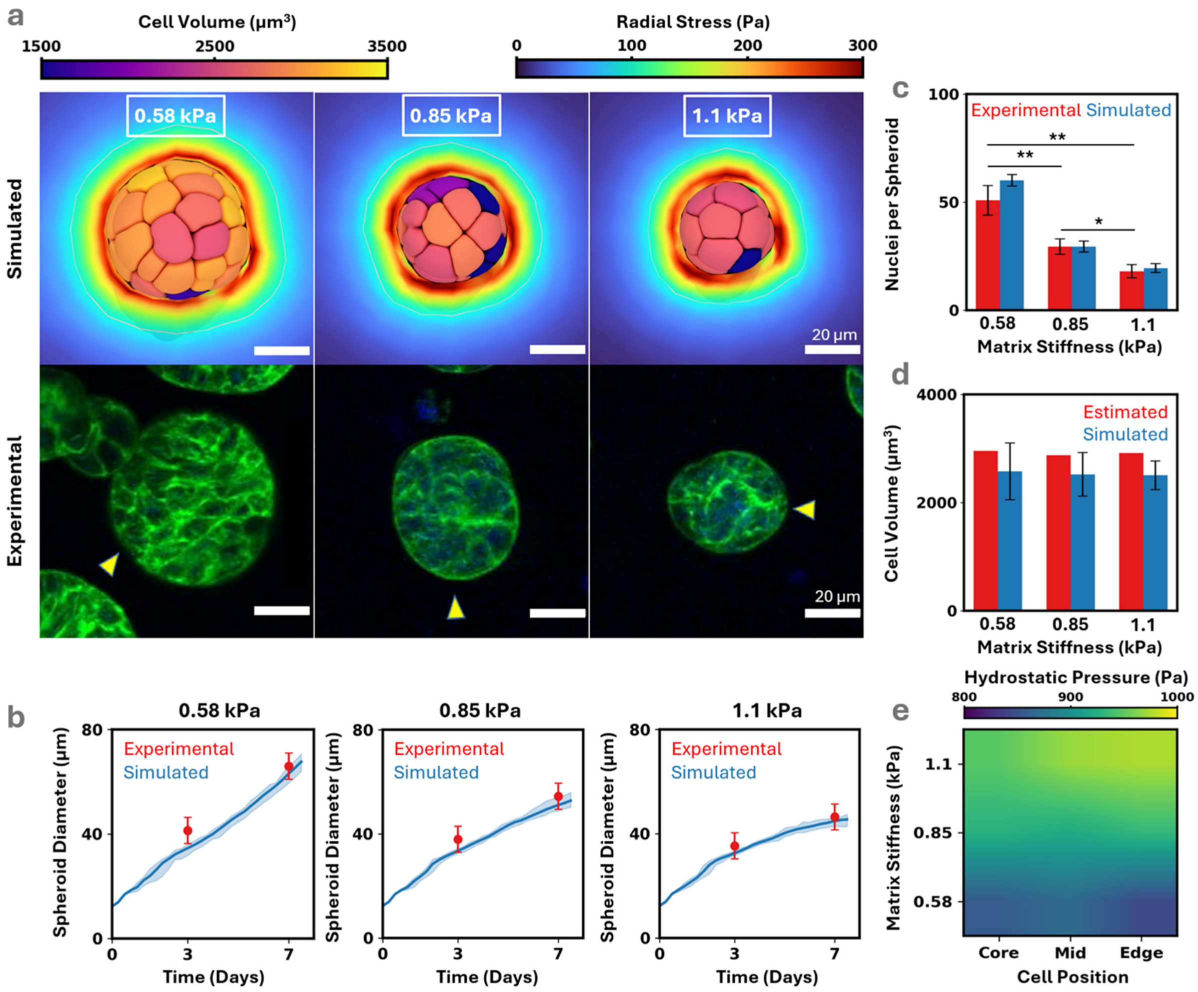


***Figure S4: 4T1 spheroid growth simulations in matrices of varying stiffness. (a)** Section cut of confined growth for three different hydrogel stiffnesses compared against experimental data. **(b)** Predicted confined spheroid growth curves closely aligns with experimental observations. The shaded blue region representing*

*the minimum and maximum spheroid diameters predicted during the simulations.* ***(c)*** *Predicted number of cells per spheroid and* ***(d)*** *cell volume align with experimental observations and estimations respectively.* ***(e)*** *The distribution of hydrostatic pressure is uniform throughout the spheroid, with hydrostatic pressure increasing with hydrogel stiffness. All error bars show mean ± SD. Significance compared using two-way ANOVA with a Bonferonni's post hoc test with *p≤0.01, ** p≤0.0001.*
